# Imaging α-synuclein pathologies in animal models and patients with Parkinson’s and related diseases

**DOI:** 10.1101/2020.10.23.349860

**Authors:** Hironobu Endo, Maiko Ono, Yuhei Takado, Kiwamu Matsuoka, Manami Takahashi, Kenji Tagai, Yuko Kataoka, Kosei Hirata, Keisuke Takahata, Chie Seki, Naomi Kokubo, Masayuki Fujinaga, Wakana Mori, Yuji Nagai, Koki Mimura, Katsushi Kumata, Tatsuya Kikuchi, Aki Shimozawa, Sushil K. Mishra, Yoshiki Yamaguchi, Hiroshi Shimizu, Akiyoshi Kakita, Hiroyuki Takuwa, Hitoshi Shinotoh, Hitoshi Shimada, Yasuyuki Kimura, Masanori Ichise, Tetsuya Suhara, Takafumi Minamimoto, Naruhiko Sahara, Kazunori Kawamura, Ming-Rong Zhang, Masato Hasegawa, Makoto Higuchi

## Abstract

Deposition of α-synuclein fibrils is implicated in Parkinson’s disease (PD) and dementia with Lewy bodies (DLB), while *in vivo* detection of α-synuclein pathologies in these illnesses has been challenging. Here, we have developed a small-molecule ligand, C05-05, for visualizing α-synuclein deposits in the brains of living subjects. *In vivo* optical and positron emission tomography (PET) imaging of mouse and marmoset models demonstrated that C05-05 captured a dynamic propagation of fibrillogenesis along neural pathways followed by disruptions of these structures. High-affinity binding of ^18^F-C05-05 to α-synuclein aggregates in human brain tissues was also proven by *in vitro* assays. Notably, PET-detectable ^18^F-C05-05 signals were intensified in the midbrains of PD and DLB patients as compared to healthy controls, providing the first demonstration of visualizing α-synuclein pathologies in these illnesses. Collectively, we propose a new imaging technology offering neuropathology-based translational assessments of PD and allied disorders towards diagnostic and therapeutic research and development.

## INTRODUCTION

Parkinson’s disease (PD) and dementia with Lewy bodies (DLB) are neurodegenerative illnesses of high prevalence with progressive motor and cognitive impairments. They are neuropathologically characterized by the appearance of Lewy bodies and Lewy neurites, which are mainly composed of aggregated α-synuclein.^1–3^ Abnormal α-synuclein is also a major component of glial cytoplasmic inclusions (GCIs) in multiple system atrophy (MSA), a neurodegenerative disease presenting with movement and autonomic disorders.^4^ In these disorders, referred to as α-synucleinopathies, ultrastructures of α-synuclein filaments containing β-pleated sheets^5^ may display diversity in disease-specific and individually variable manners as revealed by the latest cryo-electron microscopic analysis.^6^ Previous studies also experimentally demonstrated prion-like propagation of the α-synuclein fibrillogenesis throughout the brain via neural circuits.^7–9^

Formation of intracellular α-synuclein fibrils is mechanistically linked to neurodegenerative processes, and the spread of α-synuclein inclusions in the brain is supposed to be the neuropathological basis of disease progression^10–12^, supporting the significance of the α-synuclein assembly as a diagnostic and staging biomarker and a therapeutic target. In the meantime, although the possibility of detecting patients with α-synucleinopathies using cerebrospinal fluid and serum samples has been demonstrated^13,14^, the diagnosis of PD, DLB, and MSA can only become definite by examining the presence of α-synuclein aggregates in the autopsied brains. *In vivo* imaging techniques for detecting α-synuclein aggregates with high sensitivity would provide definitive information on the disease diagnosis at an early stage and could be of great utility for the evaluation of efficacies yielded by candidate drugs targeting α-synuclein pathologies at non-clinical and subsequently clinical levels.

Molecular imaging modalities, as exemplified by positron emission tomography (PET), have enabled visualization of amyloid β^15^ and tau^16–19^ deposits in the brain of living patients with Alzheimer’s disease (AD) and related disorders along with mouse models of these illnesses. A significant subset of the PET probes for these proteinopathies is a self-fluorescent β-sheet ligand and is applicable to intravital two-photon laser fluorescence microscopy of the animal models. Hence, there has been growing expectation that small-molecule ligands for β-sheet structures would also serve as PET and optical probes for multi-scale assessments of intracellular α-synuclein fibrils.^20,21^ As an initial attempt, ^11^C-BF-227, a PET ligand developed to detect amyloid β plaques^22^, was reported to bind to α-synuclein lesions in the brains of MSA patients in a PET study^23^, but *in vitro* autoradiography of postmortem MSA brain sections in a more recent study did not support significant binding of ^11^C-BF-227 to GCIs at concentrations typically achieved in PET experiments.^24^ The tau PET ligand, ^11^C-PBB3, was also documented to react with α-synuclein lesions, including Lewy bodies, Lewy neurites, and GCIs, while it has been indicated that its binding affinity for α-synuclein pathologies is not sufficient for sensitive PET detection of these lesions in living individuals.^25,26^ Indeed, PBB3 and its fluorinated analog, PM-PBB3, showed high affinity and selectivity for the β-sheet structure of tau filaments, which is assumed to be ultrastructurally distinct from that of α-synuclein assemblies.^6,19,27–31^ In the meantime, the modest binding of PBB3 and its fluorinated analog, ^18^F-PM-PBB3 (also known as florzolotau^32^), with α-synuclein inclusions implies their utility as starting compounds for the development of novel derivatives with more appropriate binding properties for *in vivo* imaging of α-synucleinopathies.

In our screening by *in vitro* evaluation, we found that derivatives of PBB3 with (*E*)-hex-2-en-4-yne linker, termed C05 series compounds, exhibited binding to α-synuclein pathologies with high reactivity and selectivity compared to PBB3 and BF-227. The *in vitro* characteristics of a chemical in this class, C05-01, were further analyzed with a tissue microarray in our earlier work.^33^ More recently, a PET imaging agent with structural similarity to the C05 series, ^18^F-SPAL-T06, enabled high-contrast imaging of α-synuclein deposits in MSA cases.^34^ PET detection of MSA α-synuclein deposits was also reported by another research group with ^18^F-ACI-12589, a radiotracer of a distinct chemotype.^35^ The success in the visualization of hallmark MSA lesions was primarily attributed to the high abundance of α-synuclein aggregates in this illness^25^, whereas ^18^F-SPAL-T06 and ^18^F-ACI-12589 have failed to capture PD and DLB pathologies. The rapid clearance of ^18^F-SPAL-T06, as compared to clinically available proteinopathy PET probes exemplified by ^18^F-PM-PBB3, allowed imaging with minimal background signals but might preclude the sustenance of free radioligand concentrations required for sufficient binding to the unabundant target molecules in Lewy bodies and neurites.

In the present study, we discovered that a C05 series compound, (*E*)-1-fluoro-3-((2-(4-(6-(methylamino)pyridine-3-yl)but-1-en-3-yn-1-yl)benzo[*d*]thiazol-6-yl)oxy)propan-2-ol (dubbed C05-05), has more suitable properties than formerly evaluated C05-01 as an *in vivo* imaging agent. Indeed, C05-05 was applicable to the detection of α-synuclein inclusions in living murine and non-human primate models of propagating α-synuclein pathologies bimodally by optical and PET imaging from single-cell to brain-wide scales, undergoing a slightly slower clearance from the brain than PM-PBB3. Furthermore, the high binding affinity of C05-05 for α-synuclein inclusions in brain tissues derived from PD, DLB, and MSA cases has supported the applicability of this probe to clinical PET in humans. Finally, exploratory clinical PET assays demonstrated the capability of ^18^F-C05-05 for detecting α-synuclein depositions in the midbrains of patients with PD and DLB.

## RESULTS

### *In vitro* binding of C05 compounds to α-synuclein pathologies

We screened PBB3 derivatives by fluorescence staining in view of the fact that most of these chemicals are self-fluorescent.^16,27^ We then identified C05 series compounds, which possess an (*E*)-hex-2-en-4-yne linker in the backbone structure, as candidates for α-synuclein imaging agents (Figure 1A). Double fluorescence staining of DLB brain slices with ligands and antibody against phosphorylated α-synuclein (pS129) demonstrated that C05-01, (*E*)-2-(4-(6-(methylamino)pyridin-3-yl)but-1-en-3-yn-1-yl)benzo[*d*]thiazol-6-ol (C05-03), and C05-05 strongly labeled Lewy bodies and Lewy neurites, whereas PBB3 and BF-227 yielded moderate and weak fluorescence signals, respectively, on these lesions (Figure 1B). To compare the binding selectivity for α-synuclein versus Aβ and tau pathologies between ligands, fluorescence staining of Lewy bodies and Lewy neurites in DLB amygdala and Aβ plaques and tau tangles in the AD middle frontal gyrus with the C05 series and reference compounds were quantified with a uniform imaging condition (Figure 1C and Figure S1). The background-corrected fluorescence intensity indicated that the signals attributed to C05-01, C05-03, and C05-05 bound to α-synuclein pathologies were significantly higher than those of Aβ and tau (Figure 1D), suggesting that the selectivity of C05 series compounds for α-synuclein versus Aβ and tau aggregates in the human brains. In contrast, the fluorescence signals originating from tau-bound PBB3 were significantly higher than those of this compound attached to α-synuclein and Aβ deposits, and the fluorescence signals attributed to Aβ-bound BF-227 were significantly higher than those of this compound attached to α-synuclein and tau deposits.

**Figure 1.**
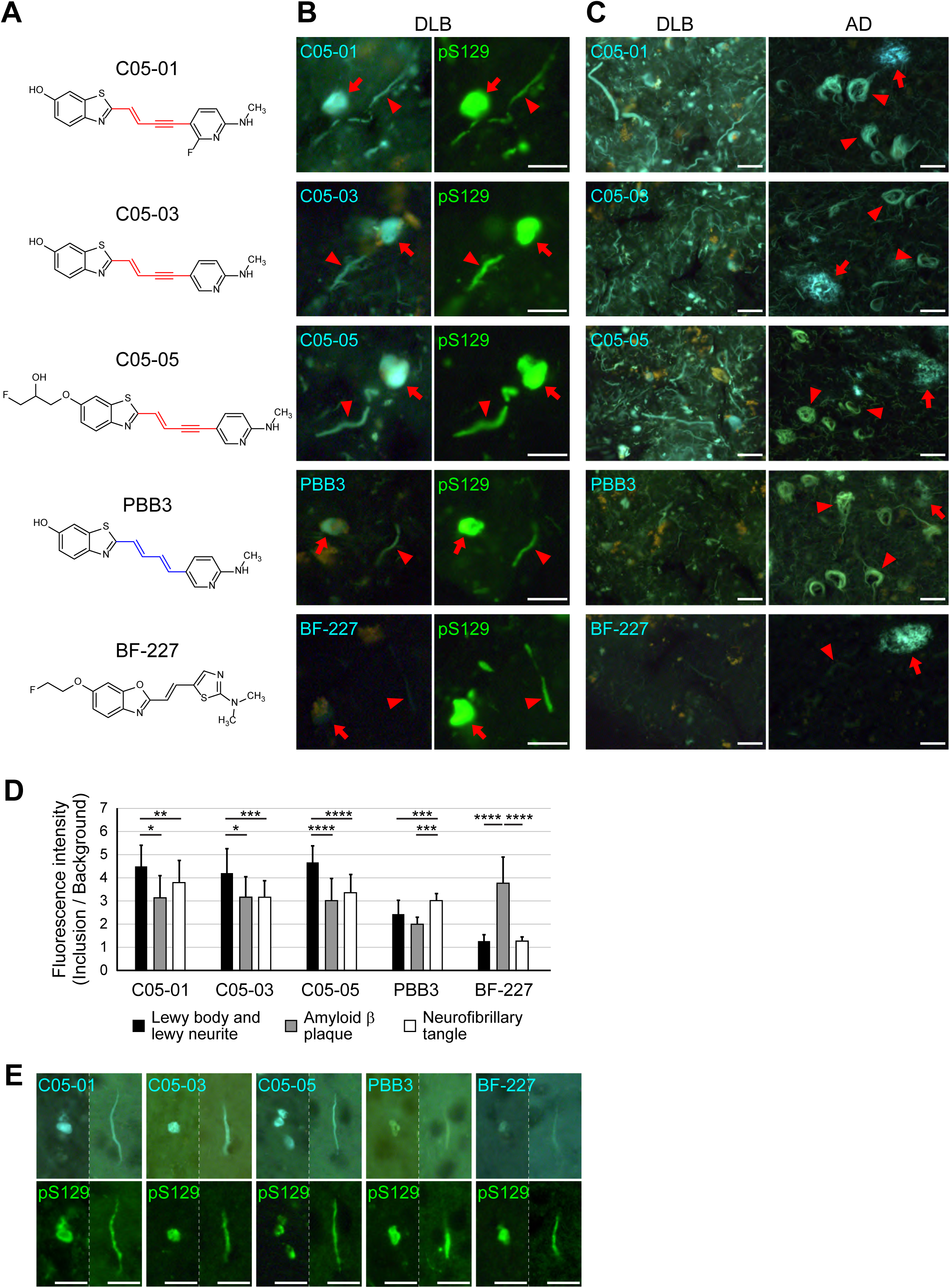
C05 series compounds bind to α-synuclein inclusions in DLB and mouse model of α-synucleinopathy *in vitro*. (**A**) Chemical structures of C05 series compounds, PBB3 and BF-227. C05-01, C05-03, and C05-05 are derivatives from PBB3 with the substitution of its (2*E*,4*E*)-hexa-2,4-diene linker (blue) with (*E*)-hex-2-en-4-yne (red). (**B**) Double fluorescence staining of Lewy bodies (arrows) and Lewy neurites (arrowheads) in the amygdala sections of a patient with DLB (also see Table S1) with 30 μM of self-fluorescent ligands (left) and anti-phosphorylated α-synuclein antibody, pS129 (right). C05-01, C05-03, and C05-05 intensely labeled α-synuclein inclusions in DLB brain sections, while PBB3 and BF-227 yielded moderate and weak staining of these lesions, respectively. (**C**) Fluorescence microscopic images of various fibrillary protein pathologies, including Lewy bodies and Lewy neurites in the amygdala sections of a patient with DLB (left) and amyloid plaques (right, arrows) and neurofibrillary tangles (right, arrowheads) in the middle frontal gyrus sections of a patient with AD (AD-1, also see Table S1), labeled with C05-01, C05-03, C05-05, PBB3, and BF-227 were taken under a uniform imaging condition. (**D**) Fluorescence signal intensities in Lewy bodies and neurites (black), amyloid plaques (gray), and neurofibrillary tangles (white) in the images illustrated in **c** were normalized according to background signals. Quantification of the background-corrected fluorescence intensity indicated that C05-01 (F(2, 74) = 6.729, *p* = 0.0021), C05-03 (F(2, 73) = 9.151, *p* = 0.0003), and C05-05 (F(2, 85) = 36.92, *p* < 0.0001) bound to α-synuclein pathologies produced significantly more intense signals than these chemicals bound to Aβ and tau pathologies. In contrast, PBB3 bound to tau pathologies elicited stronger fluorescence than this compound bound to α-synuclein and Aβ pathologies (F(2, 73) = 12.57, *p* < 0.0001), and the fluorescence signals attributed to BF-227 bound to Aβ pathologies were significantly more intense than the signals related to α-synuclein– and tau-bound BF-227 (F(2, 63) = 114.0, *p* < 0.0001). Data are presented as mean ± SD. *, *p* < 0.05; **, *p* < 0.01; ***, *p* < 0.001; ****, *p* < 0.0001 by one-way ANOVA with post-hoc Tukey’s HSD test. (**E**) Double fluorescence staining of α-synuclein inclusions resembling Lewy bodies (left) and Lewy neurites (right) in the neocortical sections of an α-Syn mouse injected with α-synuclein fibrils into the unilateral striatum (10 weeks after inoculation) with 30 μM of self-fluorescent ligands (top) and pS129 (bottom). Scale bars, 20 μm (**B**, **C**, and **E**).

### *In vivo* optical visualization of α-synuclein inclusions and tracking of pathological α-synuclein propagation with C05-05

For assessing *in vivo* detectability of individual α-synuclein inclusions by C05 series compounds, we utilized a mouse model of propagating α-synuclein fibrillogenesis induced by inoculation of recombinant mouse α-synuclein fibrils into the brain parenchyma of a wild-type mouse (α-Syn mouse).^8^ In the α-Syn mouse, aggregates of phosphorylated endogenous α-synuclein molecules emerged bilaterally in extensive brain regions, including the striatum, cortex, amygdala, and substantia nigra, from 2 weeks after unilateral inoculation of α-synuclein fibrils into the striatum (Figure S2). Double-staining of brain slices with fluorescent compounds and pS129 demonstrated that C05-01, C05-03, and C05-05 intensely labeled pS129-positive phosphorylated α-synuclein inclusions similar to Lewy bodies and Lewy neurites in the neocortex of α-Syn mice, while PBB3 and BF-227 bound modestly with these deposits (Figure 1E). We selected C05-01 and C05-05 for the following characterizations in consideration of their suitability for ^18^F radiolabeling towards broader availability.

To assess the time course of *in vivo* labeling of intraneuronal α-synuclein inclusions with C05-01 and C05-05 compared with PBB3, we conducted intravital two-photon laser fluorescence microscopy of the somatosensory cortex of an α-Syn mouse through a cranial window. Detection of C05-05, C05-01, and PBB3 signals in the same field of view of a single individual animal indicated rapid entry of C05-05 into the brain after intraperitoneal administration, reaching α-synuclein inclusions within 5 min, and the binding continued for 90 min (Figure 2A). Unlike C05-05, no noticeable increases in fluorescence signals were produced in neurons by intraperitoneally injected C05-01 and PBB3. *Ex vivo* examination of frozen brain sections from α-Syn mouse following intravital two-photon microscopy further proved the binding of intraperitoneally administered C05-05 to α-synuclein inclusions abundantly present in the somatosensory cortex of this mouse (Figure S3). These *in vivo* and *ex vivo* data collectively demonstrate the capability of C05-05 for high-contrast optical visualization of α-synuclein inclusions in a living mouse model.

**Figure 2.**
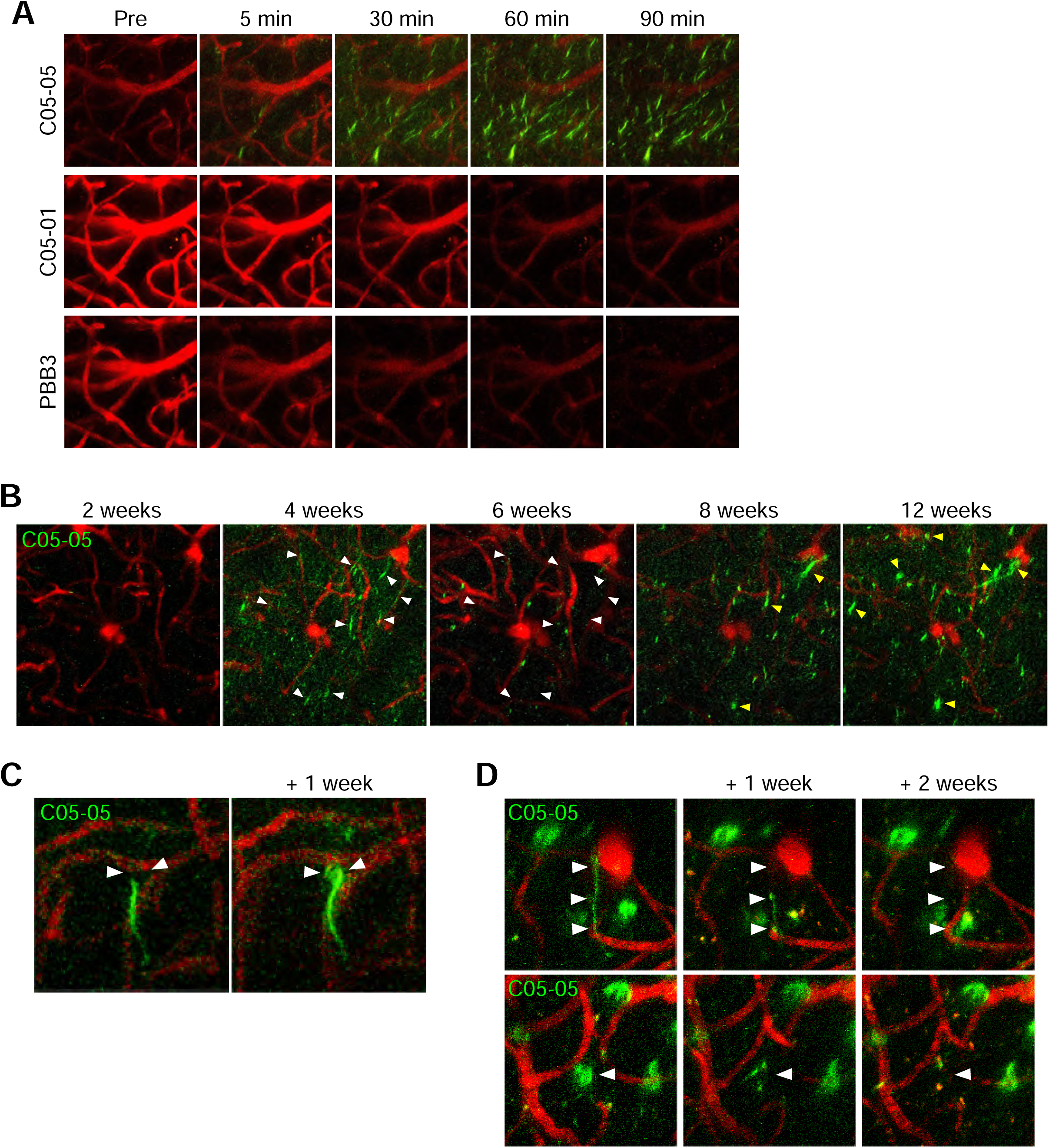
C05-05 enables *in vivo* optical visualization of individual α-synuclein inclusions in the brain of an α-Syn mouse model. (**A**) Maximum intensity projection of fluorescence signals in an identical 3D volume (field of view, 150 × 150 μm; depth, 25 – 100 μm from the brain surface) of the somatosensory cortex of a living α-Syn mouse at 8 – 10 weeks after inoculation of α-synuclein fibrils into the neocortex. Exogenous α-synuclein fibrils were found to vanish by 2 weeks after injection, followed by aggregation of endogenous α-synuclein molecules. From left, images acquired before (Pre) and 5, 30, 60 and 90 min after intraperitoneal administration of C05-05 (1.66 mg/kg) (top), C05-01 (1.66 mg/kg) (middle), and PBB3 (1.66 mg/kg) (bottom) are displayed. Somatodendritic labeling of putative neurons with C05-05 was observed as green fluorescence from 5 min after ligand administration. Fluorescence images of the corresponding area at 5 – 90 min after C05-01 and PBB3 injections demonstrated no overt retention of the tracer in the tissue. (**B**) Longitudinal *in vivo* two-photon microscopic imaging of α-synuclein inclusions with systemically administered C05-05 in the right somatosensory cortex of a single indivisual α-Syn mouse at 2, 4, 6, 8, and 12 weeks after inoculation of α-synuclein fibrils into the right striatum. A maximum projection of fluorescence in an identical 3D volume (field of view, 182 × 182 μm; depth, 40 – 400 μm from the brain surface) at 90 min after intraperitoneal administration of C05-05 demonstrated propagation of C05-05-positive α-synuclein inclusions to the cortical area from 4 weeks after the intrastriatal fibril inoculation, and subsequent changes in the subcellular location and amount of the inclusions. White arrowheads indicate neuritic α-synuclein accumulations which disappeared from 4 to 6 weeks after the fibril inoculation, and yellow arrowheads indicate somatic α-synuclein inclusions which appeared from 8 weeks after the fibril inoculation. (**C** and **D**) Longitudinal intravital microscopy of the somatosensory cortex (field of view, 55 × 55 μm; depth, 0 – 75 μm from the brain surface) of an α-Syn mouse demonstrated extension of a C05-05-positive intraneuronal α-synuclein inclusion from neurite to soma in a week (**C**, arrowheads), and disappearance of C05-05-positive neuritic inclusion similar to Lewy neurite (**D**, top, arrowheads) and somatic deposit resembling Lewy body (**D**, bottom, arrowheads) like inclusions in two weeks. Cerebral blood vessels were also labeled in red with intraperitoneally administered sulforhodamine 101.

To assess the dissemination of fibrillary α-synuclein pathologies via neuronal processes and cell bodies and consequent disruptions of these cellular structures on a longitudinal basis, we then performed biweekly intravital two-photon imaging of α-synuclein inclusions in the brain of an α-Syn mouse inoculated with α-synuclein fibrils into the unilateral striatum. C05-05-labeled neuritic α-synuclein accumulations appeared abundantly in the somatosensory cortex of the inoculated hemisphere at four weeks and decreased at six weeks (Figure 2B, white arrowheads). Moreover, the formation and growth of somatic α-synuclein inclusions labeled with C05-05 were observed eight and 12 weeks after inoculation (Figure 2B, yellow arrowheads). High-magnification images clearly visualized the intraneuronal expansion of pathological α-synuclein aggregates from neuritic to somatic compartments in a week (Figure 2C). Moreover, time-course assays provided evidence for the disappearance of neuritic and somatic α-synuclein inclusions in two weeks (Figure 2D). Our findings demonstrated the utility of C05-05 as an optical probe for a dynamic pursuit of the α-synuclein pathogenesis at a single-cell level.

### PET imaging with ^18^F-C05-05 for α-Syn mice and marmoset

The *in vivo* performance of ^18^F-labeled C05-05 (^18^F-C05-05, Figure S4) as an imaging probe was evaluated by PET scans of α-Syn mice at six months after injection of α-synuclein fibrils or saline into the bilateral striata followed by *ex vivo* autoradiography and histopathological examinations. As depicted in Figure 3A, the retention of radioligand was overtly increased in the bilateral striatal and cortical areas of an α-Syn mouse, in sharp contrast to the low radioactivity signals sustained in these brain regions of a control mouse. ^18^F-C05-05 rapidly entered the brain after intravenous administration, and peak radioactivity uptakes estimated as standardized uptake values (SUVs) were 1.19 and 1.11 in the striatum and cortex, respectively (Figure 3B, top). This was followed by a washout of radioactivity from the brain of control mice, which was slightly slower than the reported clearance of the tau PET probe, ^18^F-PM-PBB3.^36^ Relative to the controls, the radioactivity clearance was retarded in the striatum and cortex of α-Syn mice, reflecting radioligand binding to α-synuclein deposits. In the cerebellum lacking α-synuclein pathologies, there was no clear difference in the retention of radioligand between α-Syn and control mice (Figure 3A, bottom, and 3B, top), justifying the use of the cerebellum as a reference tissue for quantification of the radioligand binding. The target-to-reference ratios of the radioactivity, which is denoted as standardized uptake value ratio (SUVR), at each time point and average SUVR at 90-120 min after intravenous administration of ^18^F-C05-05 were increased in the striatum and cortex of α-Syn mice compared to those of control mice (Figure 3B, bottom).

**Figure 3.**
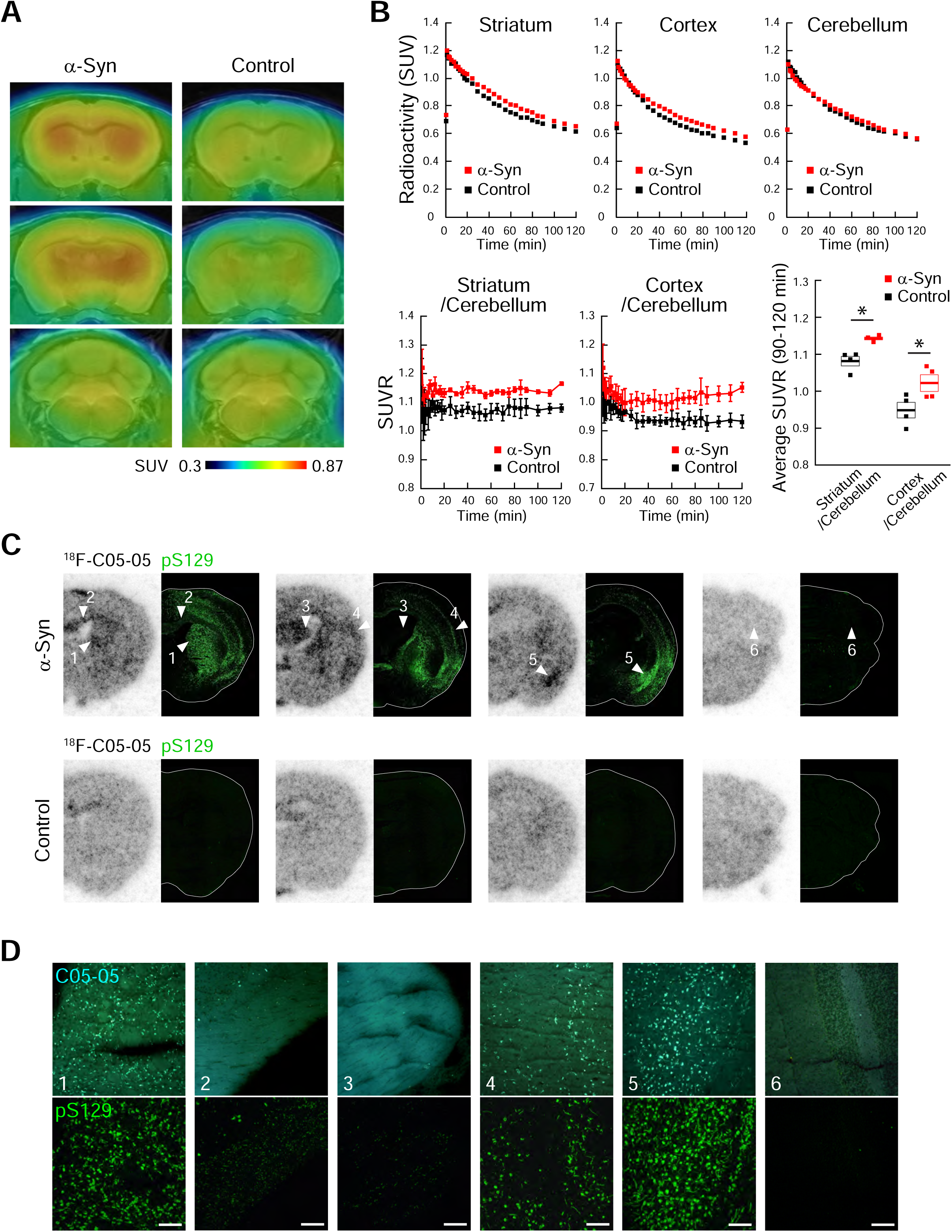
*In vivo* PET imaging with ^18^F-C05-05 detects α-synuclein deposits in the brains of α-Syn mice. (**A**) Coronal PET images at bregma +0.50 mm (top) and –0.46 mm (middle) containing the striatum and neocortex, and –6.64 mm (bottom) containing the cerebellum generated by averaging dynamic scan data at 60 – 90 min after intravenous administration of ^18^F-C05-05 (30.8 ± 0.4 MBq) in mice at 6 months after inoculation of α-synuclein fibrils (α-Syn mouse, left) or saline (control mouse, right) into the bilateral striata. PET images are superimposed on a magnetic resonance (MR) image template. Voxel values represent SUV. (**B**) Time-radioactivity curves in the striatum, neocortex, and cerebellum during the dynamic PET scan (top), time-course changes in the target-to-cerebellum ratio of radioactivity (SUVR, left and middle panels in bottom row), and the average of target-to-cerebellum ratios at 90 – 120 min (bottom, right) in α-Syn (red symbols) and control (black symbols) mice. There were significant main effects of animal group and region in two-way, repeated-measures ANOVA (group, F(1, 6) = 11.39, *p* = 0.015; region, F(1, 6) = 111.9, *p* < 0.0001). *, *p* < 0.05 by Bonferroni’s post hoc test. Data are presented as mean (top) or mean ± SEM (bottom) in four α-Syn or control mice. (**C**) *Ex vivo* examination of frozen brain sections obtained from α-Syn (top) and control (bottom) mice after PET imaging to assess distributions of intravenously administered ^18^F-C05-05 (27.8 ± 0.2 MBq), in comparison with immunolabeling of the same sections with pS129. From left, coronal brain sections at bregma +0.50, –0.46, –1.94, and –6.64 mm are displayed. (**D**) High-power photomicrographs showing double fluorescence staining of the section used for *ex vivo* examination with 30 μM of C05-05 (top) and pS129 (bottom). Areas correspond to those indicated by arrowheads in C. The striatum (**1**), somatosensory cortex (**4**), and amygdala (**5**) of an α-Syn mouse contained abundant α-synuclein inclusions. The corpus callosum (**2**) and fimbria of the hippocampus (**3**) showed a small number of α-synuclein deposits. The cerebellum (**6**) contained very few α-synuclein inclusions. Scale bars, 100 μm.

*Ex vivo* autoradiography of brain tissues collected from α-Syn and control mice used for PET scan at 90 min after intravenous ^18^F-C05-05 injection demonstrated accumulation of the radioligand in the striatum, cortex, and amygdala of the α-Syn mouse harboring abundant neuronal α-synuclein inclusions (Figure 3C and D). Conversely, there was no noticeable increase of radioligand retentions in these brain regions of the control mouse. In addition, the radioligand accumulation was minimal in the cerebellum of the α-Syn and control mice, the area devoid of α-synuclein deposits, while non-specific radioligand accumulations in several white matter regions, including the corpus callosum and fimbria of the hippocampus, was observed in both of these mice.

Since the small brain volumes of mice impeded clear separations between striatal and neocortical radioactivity signals as assessed by PET, the *in vivo* traceability of the inter-regional α-synuclein dissemination with the use of ^18^F-C05-05 remained rather inconclusive. We accordingly employed a non-human primate model of propagating α-synuclein pathologies by inoculating recombinant marmoset α-synuclein fibrils into the brain parenchyma (α-Syn marmoset).^37^ An α-Syn marmoset receiving marmoset α-synuclein fibrils exhibited enhanced retention of ^18^F-C05-05 in a sub-portion of the caudate nucleus containing the injection site at one month after inoculation, which spread extensively in the caudate nucleus, putamen, and substantia nigra of the ipsilateral hemisphere and to a lesser extent in the contralateral left hemisphere at three months (Figure 4A). We also performed PET imaging of dopamine transporters with a specific radioligand, ^11^C-PE2I, which has proven useful for detecting degenerations of dopaminergic neurons in PD and its models^38,39^, in the α-Syn marmoset before (Pre) and three months after inoculation. Parametric images of ^11^C-PE2I binding potential (BP_ND_) demonstrated a decrease of dopamine transporters in the caudate nucleus, putamen, and substantia nigra of the inoculated hemisphere compared to the contralateral hemisphere (Figure 4B), in agreement with the distribution of augmented ^18^F-C05-05 retentions and reported pathological α-synuclein depositions.^37^

**Figure 4.**
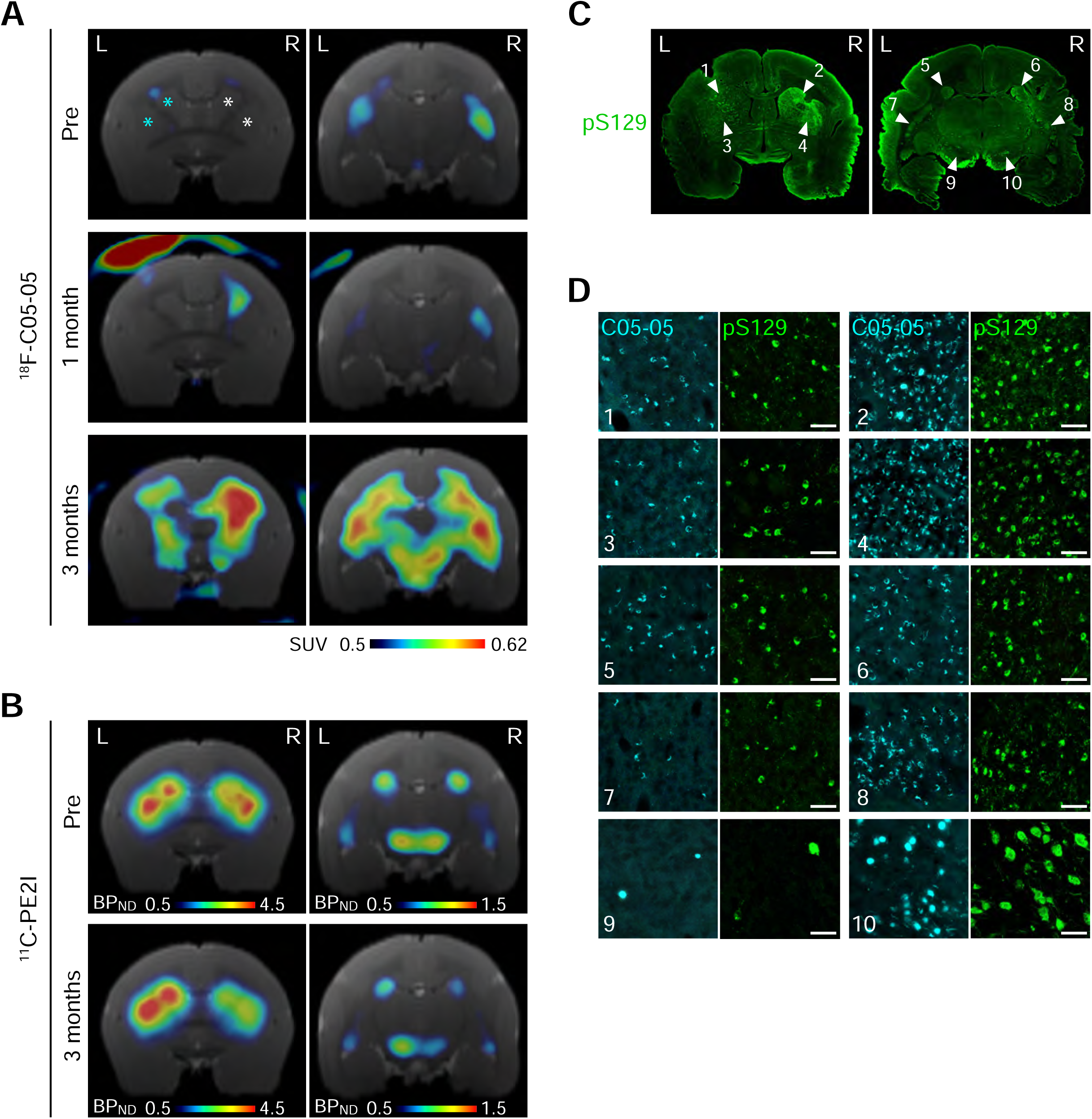
Longitudinal *in vivo* PET imaging with ^18^F-C05-05 visualizes the propagation of pathological α-synuclein aggregates in the brain of an α-Syn marmoset. (**A**) Coronal brain images in a marmoset injected with α-synuclein fibrils and saline into the right and left caudate nucleus and putamen, respectively, generated by averaging dynamic PET data at 30 – 120 min after intravenous administration of ^18^F-C05-05 (89.6 ± 15.3 MBq) (also see Figure S14). Images were acquired before (Pre), and 1 and 3 months after the fibril inoculation, and white and blue asterisks indicate the sites of α-synuclein fibril and saline injections, respectively. Brain volume data were sectioned at 9.5 mm (left) and 5.0 mm (right) anterior to the interaural line to generate images containing the caudate nucleus/putamen and caudate nucleus/putamen/substantia nigra, respectively. PET images are superimposed on an MR image template, and voxel values represent SUV. Longitudinal ^18^F-C05-05-PET showed the expansion of radioactivity signals from a part of the right caudate nucleus to extensive brain areas, including bilateral regions of the caudate nucleus, putamen, and substantia nigra from 1 to 3 months after inoculation. (**B**) Parametric images of BPND for ^11^C-PE2I (radioactivity dose: 89.2 ± 2.0 MBq) in a single individual α-Syn marmoset demonstrated reduction of the radioligand binding in the right caudate nucleus, putamen, and substantia nigra at 3 months after inoculation compared to the baseline before inoculation (Pre). Brain volume data were sectioned at 9.5 mm (left) and 5.0 mm (right) anterior to the interaural line, and BPND images were superimposed on an MR image template. (**C**) Histopathological assays were carried out 1 month after the final PET scan, demonstrating a similarity between the regional distributions of α-synuclein inclusions stained with pS129 and localization of radioligand retentions in ^18^F-C05-05-PET images at 3 months. (**D**) High-power photomicrographs showing fluorescence staining of brain sections shown in B with pS129 and adjacent brain sections with 30 μM of C05-05. Areas correspond to those indicated by arrowheads in **b**. The right caudate nucleus (**2** and **6**), putamen (**4** and **8**), and substantia nigra (**10**) contained highly abundant α-synuclein inclusions. The left caudate nucleus (**1** and **5**) and putamen (**3** and **7**) contained moderate amounts of α-synuclein deposits, and the left substantia nigra (**9**) contained sparse α-synuclein inclusions. Scale bars, 50 μm.

The brain of this animal was sampled at four months after inoculation, and immunohistochemical analyses of the brain slices demonstrated the distribution of pS129-stained α-synuclein inclusions in agreement with *in vivo* PET findings with ^18^F-C05-05 at three months (Figure 4C). Double-staining with C05-05 and pS129 confirmed dense accumulations of α-synuclein aggregates in neuronal processes and somas recapitulating PD and DLB pathologies in the caudate nucleus, putamen, and substantia nigra of the inoculated hemisphere (Figure 4D). The corresponding brain areas of the contralateral hemisphere contained less abundant α-synuclein inclusions in neurites and neuronal somas. These α-synuclein pathologies were fluorescently labeled with C05-05, suggesting that the increased retention of radioligand in ^18^F-C05-05-PET stemmed from its *in vivo* interaction with α-synuclein inclusions. Meanwhile, non-specific accumulations of radioligand in bilateral white matter regions flanking the putamen were noted in the pre-inoculation ^18^F-C05-05-PET (Figure 4A, top), and the absence of α-synuclein deposits in these areas was ensured by histochemical and immunohistochemical assays.

These *in vivo* data provide the first PET demonstration of time-course imaging of pathological α-synuclein deposits in living animal models along the course of spatially expanding fibrillogenesis accompanied by the degeneration of neural circuits involved as dissemination pathways.

### Binding of ^18^F-C05-05 to human α-synuclein pathologies

To assess the binding of ^18^F-C05-05 to human α-synuclein pathologies at a low concentration, we performed *in vitro* autoradiography of basal ganglia from MSA and Parkinson’s disease with dementia (PDD) cases and amygdala from a DLB case (Figure 5A). The total binding of ^18^F-C05-05 was markedly abolished by excessive non-radiolabeled C05-05, indicating the saturability of the radioligand binding. The MSA cases showed specific binding of ^18^F-C05-05 in association with the local α-synuclein burden (Figure 5A and B). A case with mild pathology (MSA-1) had no binding of ^18^F-C05-05 to the striatopallidal fibers. MSA-2, burdened with moderate α-synuclein deposits, showed weak ^18^F-C05-05 radioactivity signals in striatopallidal fibers containing numerous GCIs. MSA-3, which is a case with severe α-synuclein pathologies, exhibited intense radioligand binding to the striatopallidal fibers harboring densely packed GCIs. The DLB and PDD cases also showed specific binding of ^18^F-C05-05 in line with the distribution of α-synuclein pathology in the amygdala and substantia nigra, respectively.

**Figure 5.**
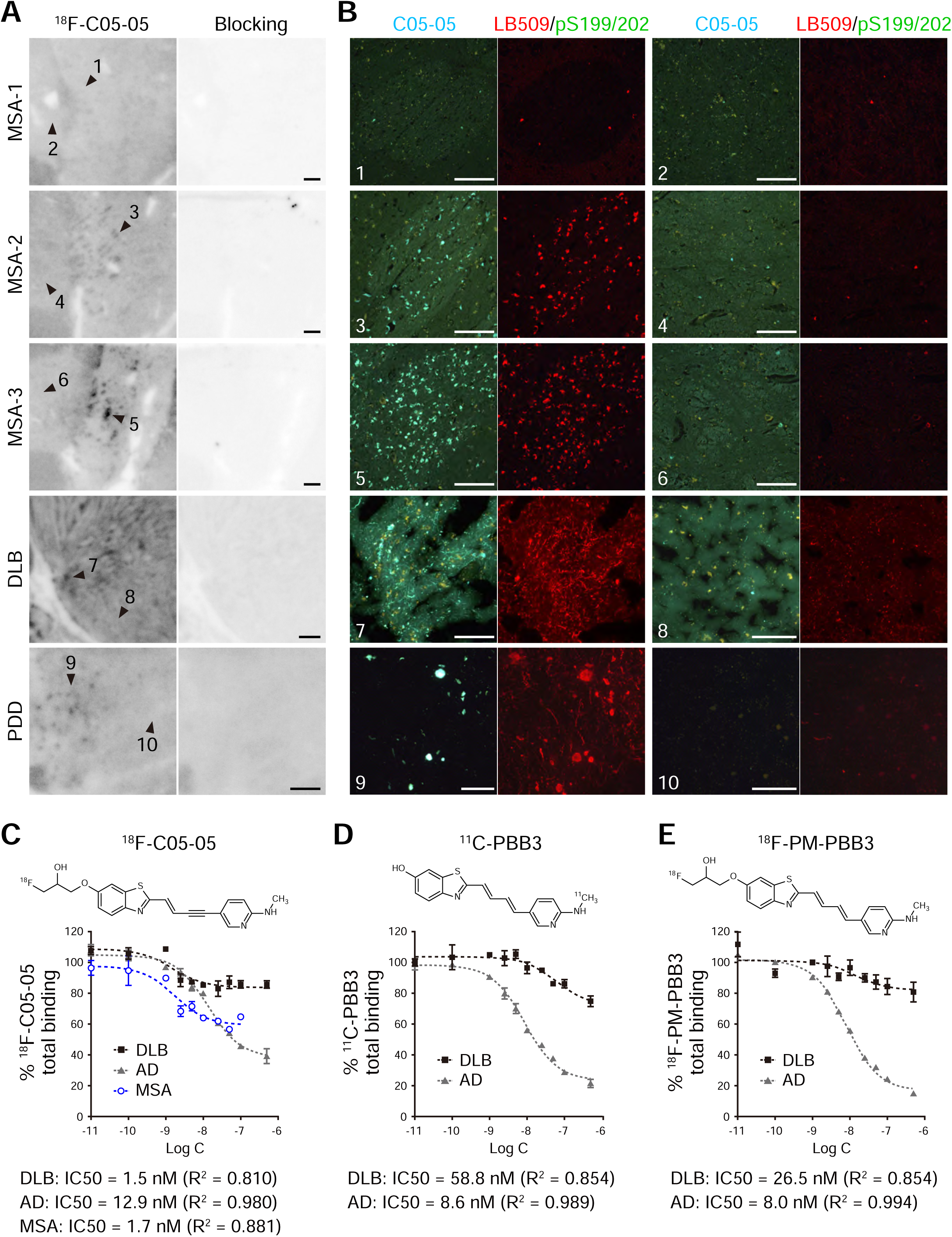
^18^F-C05-05 displays high-affinity binding to α-synuclein pathologies in DLB, MSA and PD brain tissues. (**A** and **B**) Autoradiographic labeling of sections, including the basal ganglia derived from patients with MSA (MSA-1, 2, and 3, also see Table S1), amygdala derived from patients with DLB, and substantia nigra derived from patients with PDD (also see Table S1), with 10 nM of ^18^F-C05-05 in the absence (**A**, left) and presence (**A**, right) of 10 μM of non-radiolabeled C05-05, and high-power photomicrographs showing triple fluorescence staining of the section used for ^18^F-C05-05 autoradiography with 30 μM of C05-05, LB509, and pS199/202 (**B**). Areas in **B** correspond to locations indicated by arrowheads in **A**. No overt specific binding of ^18^F-C05-05 was detected in the striatopallidal fibers (**1**) of MSA-1 with mild pathology, weak but clearly noticeable radioligand binding to these fibers (**3**) was seen in MSA-2 with moderate pathology, and strong radioligand binding to the same subregion (**5**) was observed in MSA-3 with severe pathology. No significant binding of ^18^F-C05-05 was shown in the areas devoid of α-synuclein pathologies in MSA cases (**2**, **4**, and **6**). In the amygdala of a DLB case and the substantia nigra of a PDD case, binding of ^18^F-C05-05 was seen in an area harboring abundant Lewy bodies and Lewy neurites (**7** and **9**). In contrast, no significant binding of ^18^F-C05-05 was noted in an area with a very small amount of α-synuclein pathologies (**8** and **10**). Immunohistochemistry with pS199/202 indicated the absence of tau deposits in these regions. Scale bars, 1 mm (**A**) or 100 μm (**B**). (**C**-**E**) Total (specific + non-specific) binding of ^18^F-C05-05 (**C**), ^11^C-PBB3 (**D**), and ^18^F-PM-PBB3 (**E**) in the DLB amygdala (black squares, also see Table S1) and AD frontal cortex (grey triangles, AD-2, also see Table S1) samples homologously blocked by non-radiolabeled C05-05, PBB3, and PM-PBB3, respectively, with varying concentrations. Total binding of ^18^F-C05-05 in the MSA putamen sample homologously blocked by non-radiolabeled C05-05 with varying concentrations was indicated in **C** (blue circles, also see Table S1). Data are mean ± SD in four samples and are expressed as % of average total binding. Homologous blockades of ^18^F-C05-05, ^11^C-PBB3, and ^18^F-PM-PBB3 binding described by a one-site model and parameters resulting from curve fits.

We then conducted triple staining of the sections used for autoradiography with C05-05 and antibodies against α-synuclein (LB509) and phosphorylated tau (pS199/202). The fluorescence labeling with C05-05 and LB509 was noted on GCIs in the MSA striatopallidal fibers, and Lewy bodies and Lewy neurites in the DLB amygdala and PDD substantia nigra, and these areas were devoid of pS199/202-immunoreactive phosphorylated tau pathologies (Figure 5B).

We also quantified the affinity of ^18^F-C05-05 for α-synuclein aggregates in homogenized DLB amygdala tissues in comparison to AD frontal cortical tissues. Radioligand binding in these tissues was homologously blocked by non-radiolabeled C05-05 in a concentration-dependent fashion (Figure 5C), indicating binding saturability. ^18^F-C05-05 displayed high-affinity binding in DLB homogenates with the concentration inducing 50% homologous inhibition (IC50) of 1.5 nM. This radioligand was not highly binding with Aβ and tau aggregates in AD tissues relative to DLB α-synuclein deposits, with IC50 of 12.9 nM. Unlike ^18^F-C05-05, tau PET tracers, ^11^C-PBB3 and ^18^F-PM-PBB3, displayed relatively low affinities for α-synuclein deposits in DLB homogenates with IC50 values of 58.8 nM and 26.5 nM, respectively, while these radioligands more tightly bound to AD-type protein fibrils than α-synuclein aggregates, with IC50 values of 8.6 nM and 8.0 nM, respectively (Figure 5D and E). The molecular docking analyses based on the cryo-EM structure also suggested that C05-05 binds to the potential binding sites in MSA-type α-synuclein filaments with a stronger affinity than PBB3 and PM-PBB3 (Figure S5). In fact, ^18^F-C05-05 displayed high-affinity binding in MSA homogenates with IC50 values of 1.7 nM (Figure 5C), which was comparable to the affinity of ^18^F-SPAL-T-06 (KD, 2.5 nM^34^). These results of *in vitro* autoradiographic and radioligand binding assays highlight the reactivity of ^18^F-C05-05 with human α-synuclein pathologies with much higher affinity than existing PET tracer for non-α-synuclein aggregates, supporting the potential of this novel radioligand for visualizing hallmark lesions in living patients with α-synucleinopathies.

### PET detection of α-synuclein pathologies in MSA, PD, and DLB patients with ^18^F-C05-05

Since the non-clinical results supported the utility of ^18^F-C05-05 as a sensitive α-synuclein imaging probe with high affinity and considerably long retention of free and specifically bound compounds in the brain, we conducted an exploratory clinical PET study primarily targeting α-synuclein depositions in PD and DLB, collectively referred to as Lewy body diseases (LBDs). Five patients meeting clinical diagnostic criteria for LBDs and eight healthy controls (Table 1) underwent PET imaging with ^18^F-C05-05. ^18^F-C05-05 uptake peaked rapidly after radioligand injection and then decreased by about 50% or more in most regions over the next 50 min, resulting in uniformly low radioligand retention (Figure S6). Similar to murine PET observations, the clearance of ^18^F-C05-05 from the brain was slower than the reported washout of ^18^F-PM-PBB3.^19^ Radiochromatographic data indicated progressive increases of radiometabolites in plasma, which appeared more polar than the parent molecule according to the reverse-phase HPLC retention times and are, therefore, unlikely to efficiently enter the brain (Figure S6A, B).

**Table 1.** Demographics of individuals with HCs and LBDs included in the present study.

|  | HCs | LBDs | *P* values |
| --- | --- | --- | --- |
| Number (race: all Asian) | 8 | 5<br>(PD = 4, DLB = 1) |  |
| Age | 66.1 ± 6.2 | 67.8 ± 3.4 | 0.54 |
| Female N [%] | 4 [50] | 1 [20] | 0.002 |
| Amyloid $\beta$ positive | 0 | 0 | |
| Disease duration (year) |  | 6.8 ± 3.7 |  |
| UPDRS I | N.E. | 10.6 ± 6.3 |  |
| UPDRS II | N.E. | 11.4 ± 9.5 |  |
| UPDRS III | 1.6 ± 1.9 | 39.6 ± 14.2 | 0.002 |
| MMSE | 29.5 ± 0.8 | 29.6 ± 0.6 | >0.99 |
| FAB | 16.9 ± 1.6 | 17.0 ± 1.2 | 0.84 |
| GDS | 1.5 ± 1.5 | 5.2 ± 3.6 | 0.09 |
| Apathy | 8.5 ± 5.7 | 14.8 ± 4.0 | 0.10 |

Abbreviations: FAB; Frontal Assessment Battery, GDS; Geriatric Depression Scale, HCs; Healthy Controls, LBDs; patients with Lewy body disease, MMSE; Mini-Mental State Examination, N; number, N.E.; Not Examined, PD; Parkinson's Disease, DLB; Dementia with Lewy Body, UPDRS I, II, III; Unified Parkinson's Disease Rating Scale part I, II and III.

We focused on the detection of α-synuclein pathologies in the midbrain, as the midbrain substantia nigra is burdened with Lewy pathologies commonly in ‘body-first’ and ‘brain-first’ subtypes of LBD at a clinical stage.^40^ It was documented that α-synuclein deposits were present in cerebellum of LBD cases^41,42^, which may impede the use of this anatomical structure as reference tissue for the quantification of the radioligand binding. Hence, we defined a reference region in deep white matter as in previous analyses (See STAR★METHODS).^43^ As the radioactivity concentrations in the midbrain and deep white matter of healthy controls became almost equivalent at 100 min (Figure S6C), we assessed the radioligand accumulation by quantifying SUVR at 100 – 120 min. Unlike healthy controls, LBD patients presented higher radioactivity concentration in the midbrain than deep white matter (Figure S6C). Correspondingly, parametric PET images showing SUVR at 100 – 120 min demonstrated that the retention of ^18^F-C05-05 in the midbrain of LBD cases was enhanced as compared to controls (Figures 6A and S7).

**Figure 6.**
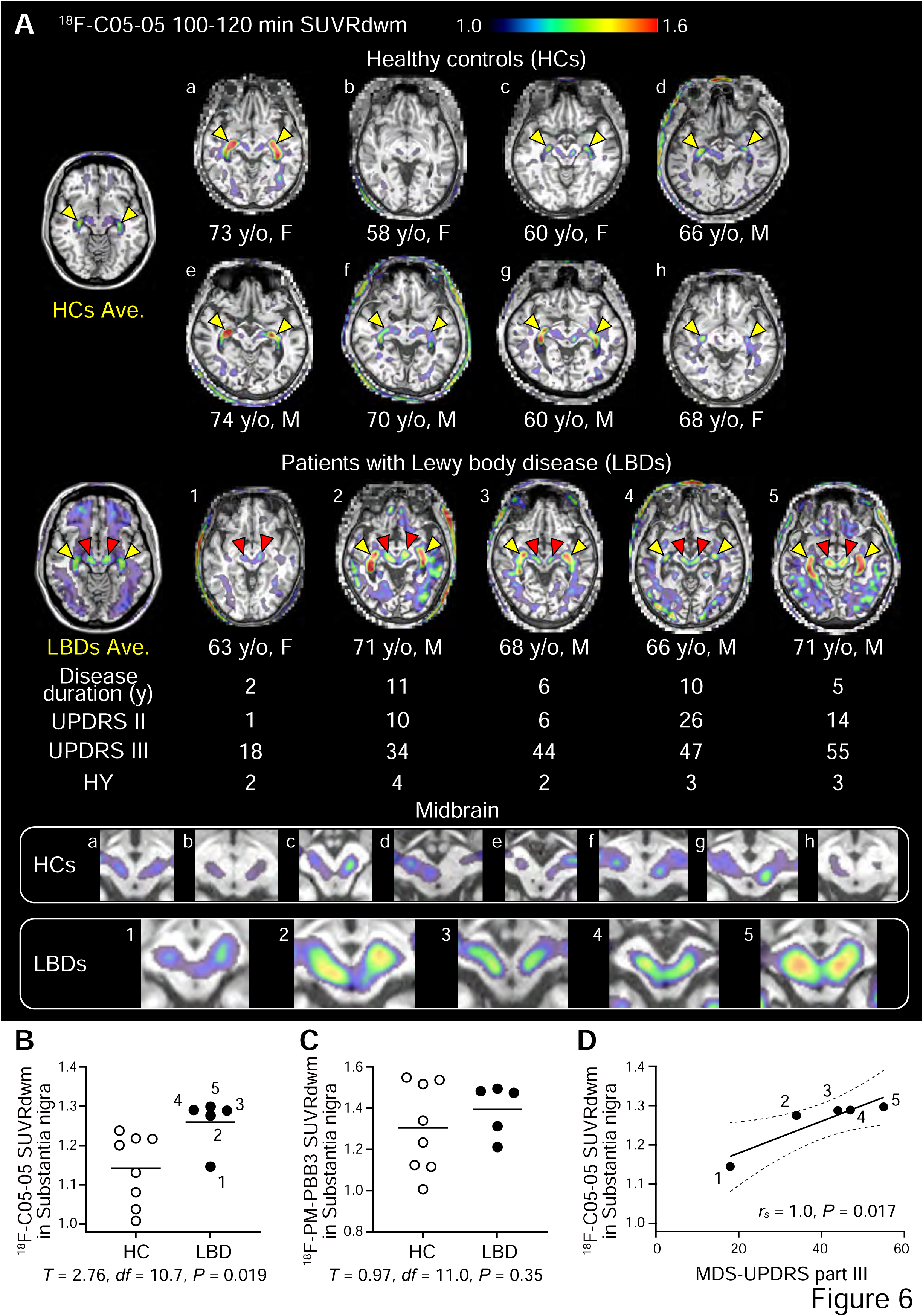
Detection of α-synuclein pathologies in the midbrains of LBD patients by PET with ^18^F-C05-05. (**A**) Parametric images of SUVRs at 100 – 120 min after ^18^F-C05-05 injection created by axial sectioning at the midbrain level with the deep white matter as the reference region (SUVRdwm). (**B, C**) Group comparison of SUVR values for ^18^F-C05-05 (**B**) and ^18^F-PM-PBB3 (**C**) in the substantia nigra between HCs and LBD patients. (**D**) Correlation between nigral SUVR values for ^18^F-C05-05 and the MDS-UPDRS (Movement Disorder Society revised Unified Parkinson’s Disease Rating Scale) part III scores in LBD cases. Spearman’s rho correlation coefficient (*rs*) is displayed in the graph. Age (years old; y/o) and sex (F, female; M, male) of the subjects are indicated, and images on the far left are averages of spatially normalized images derived from eight HCs (HCs Ave.) and five LBD cases (LBDs Ave.) in the Montreal National Institute (MNI) space (midbrain substantia nigra level corresponding to Z = –11). High-magnification images of the midbrains of subjects a – h in the HC group and 1 – 5 in the LBD group are displayed at the bottom. Red arrowheads indicate specific radioactivity accumulations in the substantia nigra, and nonspecific accumulations in the choroid plexus are denoted by yellow arrowheads. Abbreviations: Ave. (Averaged image), HY (Hoehn and Yahr stage), UPDRS II (Unified Parkinson’s Disease Rating Scale part II), UPDRS III (part III), y: (year).

These subjects also received PET scans with ^18^F-PM-PBB3 and ^11^C-PiB to visualize tau and amyloid β lesions. They were all negative for amyloid β pathologies and exhibited no accumulation of ^18^F-PM-PBB3 in the midbrain (Figures S8 and S9). Group comparisons between LBD and control subjects also showed markedly elevated SUVRs for ^18^F-C05-05 but not ^18^F-PM-PBB3 in the substantia nigra (Figure 6B, C). These observations ruled out the presence of tau aggregates in the brainstem and indicated the incapability of ^18^F-PM-PBB3 for capturing Lewy pathologies in the nigral area, which is in line with our previous finding.^19^ Despite a small sample size, there was a significant correlation between the nigral SUVR value and the degree of motor impairments assessed by Movement Disorder Society revised Unified Parkinson’s Disease Rating Scale part III scores (Figure 6D, S7). We also found moderately increased retentions of ^18^F-C05-05 in the middle temporal lobe (Figure 6), thalamus, and globus pallidus (Figure S8) of several LBD cases. Thus, the present clinical dataset documents the first successful *in vivo* visualization of α-synuclein pathologies in patients with PD and DLB.

To evaluate the detectability of α-synuclein accumulations in the MSA brains, ^18^F-C05-05 PET scans were performed in patients meeting clinical diagnostic criteria for MSA with predominant parkinsonism (MSA-P; Table S2). These individuals showed higher retentions of ^18^F-C05-05 in the putamen (Figure 7), in agreement with the *in vitro* autoradiographic data (Figure S10). In addition, two of these cases presented mild cerebellar ataxia and high radioligand accumulation in the middle cerebellar peduncle (Figure 7). These results further justify the utilization of ^18^F-C05-05 as a PET probe possessing adequate reactivity with α-synuclein aggregates in humans.

**Figure 7.**
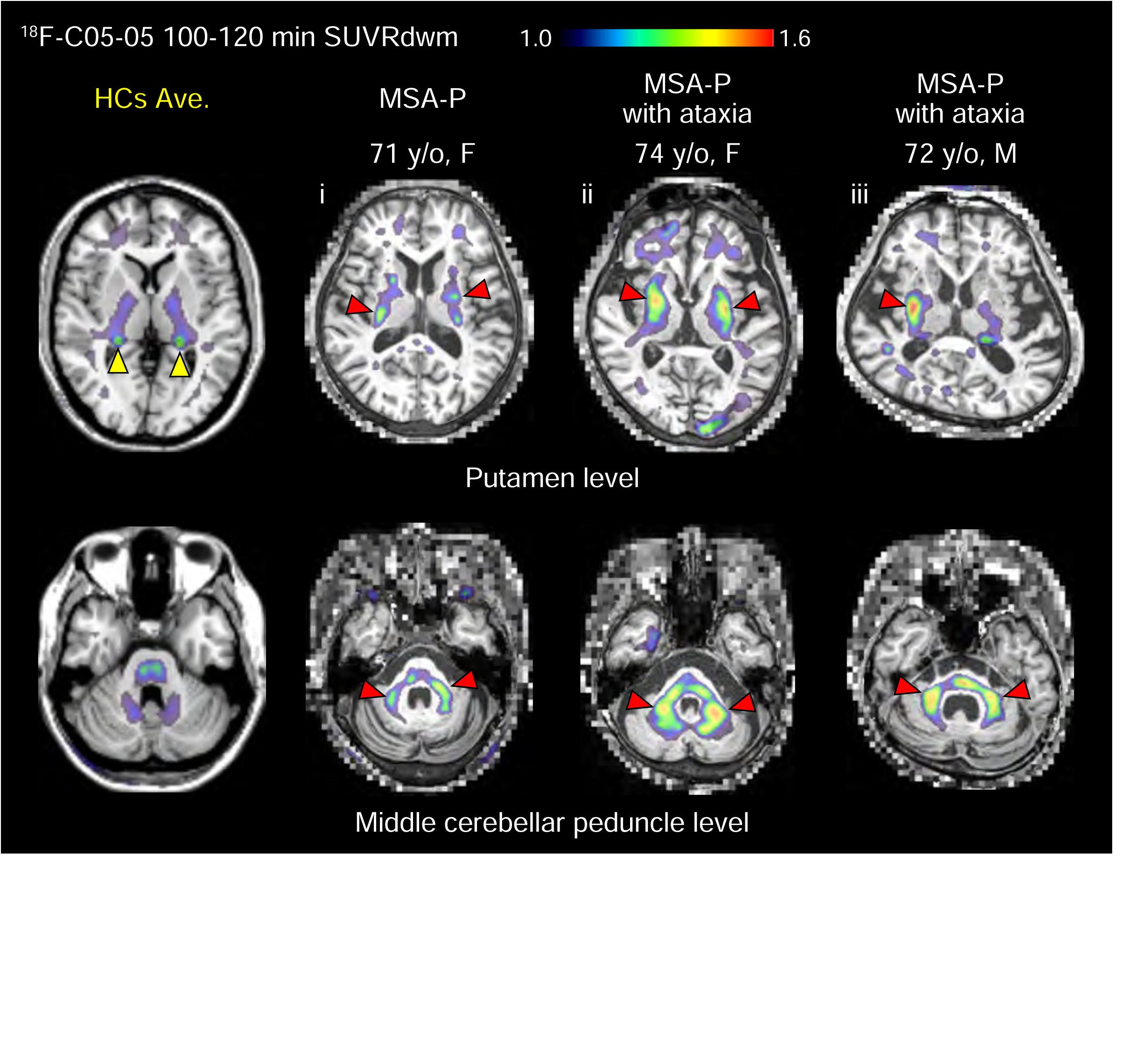
PET imaging of α-synuclein pathologies in MSA-P patients with ^18^F-C05-05. Parametric images of SUVR at 100 – 120 min after ^18^F-C05-05 injection created by axial sectioning at levels of the putamen (top row) and middle cerebellar peduncle (bottom row) with the deep white matter as the reference region are shown. Images on the far left are averages of spatially normalized images derived from eight HCs (HCs Ave.) in the Montreal National Institute (MNI) space (middle cerebellar peduncle level corresponding to Z = –30 and putamen level corresponding to Z = 3). Age (years old; y/o) and sex (F, female; M, male) of the subjects are indicated. There were accumulations of ^18^F-C05-05 radiosignals in the lateral putamen of all MSA-P cases and the middle cerebellar peduncle in MSA-P cases with mild cerebellar ataxia (red arrowheads), and nonspecific radioactivity accumulations in the choroid plexus (yellow arrowheads).

## Discussion

The current work has offered a powerful imaging tool to pursue the molecular and cellular mechanisms of the neurodegenerative α-synucleinopathies in basic research on animal models, and this technology is translated to clinical PET assessments of LBDs. The imaging methodology with ^18^F-C05-05 opens an avenue towards the neuropathology-based diagnosis and differentiation of these disorders, along with the discovery and development of anti-α-synuclein therapeutics as disease-modifying treatments.^44,45^ Our bimodal *in vivo* optical and PET assays of model animals have allowed longitudinal tracking of α-synuclein propagations through neural pathways from subcellular to brain-wide scales, facilitating the non-clinical evaluation of efficacies exerted by a candidate drug counteracting the etiological processes of α-synucleinopathies. The same PET assay has proven capable of visualizing core lesions residing in the midbrains of PD and DLB cases for the first time, besides the basal ganglia and deep cerebellar structures of MSA subjects.

It is noteworthy that the substitution of the (*2E*, *4E*)-hexa-2, 4-dien linker in the chemical structure of a tau imaging agent, PBB3, with (*E*)-hex-2-en-4-yne resulted in a profound increase of the ligand binding to α-synuclein versus tau and Aβ fibrils, leading to the generation of C05 series compounds. The molecular docking analyses based on the cryo-EM structure suggested that PBB3 and C05-05 bind to AD-type tau filaments and MSA-type α-synuclein filaments, respectively, in a direction perpendicular to the fibril axis (Figure S5).^29^ The linker substitution could produce differences in the backbone twist angle between PBB3 and C05 series compounds at a minimum energy state, which may affect the fitness of the chemical for the binding surface on the filament. Our molecular docking analysis suggested that C05-05 reacts with the potential binding sites in MSA-type α-synuclein filaments with a stronger affinity than PBB3 and PM-PBB3. In fact, IC50 of C05-05 for the homologous binding blockade in homogenized DLB amygdala tissues was approximately 40– and 18-fold smaller than those of PBB3 and PM-PBB3, respectively, implying the critical role of the linker angle in the ligand affinity for pathological fibrils. It is yet to be clarified whether C05 series compounds exhibit differential reactivity with PD, DLB, and MSA α-synuclein fibrils, although autoradiographic labeling of pathological inclusions in those illnesses was demonstrated with ^18^F-C05-05 in the present assay.

Intravital two-photon laser microscopy with C05-05 has enabled longitudinal imaging of the α-synuclein fibrillogenesis at a subcellular scale in the brain of a living α-synucleinopathy mouse model, visualizing the dynamic processes in the formation of α-synuclein lesions, including a spatiotemporal connection between the developments of Lewy neurite-like neuritic aggregates and Lewy body-like somatic inclusion in a single neuron, as well as the disappearance of these fibrillar deposits. While the vanishment of α-synuclein fibrils may reflect the loss of neurons loaded with inclusions, the dynamic appearance and disappearance of the aggregates may also unfold as a consequence of continuous translocations of α-synuclein assemblies through neuritic processes. The mechanisms linking the accumulations of α-synuclein fibrils and loss of neurons or their substructures remain elusive, and cell-autonomous^46^ and non-cell-autonomous^47^ death of neurons could be provoked in the pathogenetic pathway. Such etiological cellular events will be microscopically examined by monitoring interactions between neuronal and glial cells expressing fluorescent proteins and C05-05-positive α-synuclein deposits.

Since our longitudinal PET scans with ^18^F-C05-05 have successfully captured the dissemination of α-synuclein pathologies in an α-Syn marmoset along the course following the fibril inoculation, this imaging technology will pave the way to the neuroimaging-based evaluations of the disease severity and progression in α-synucleinopathy patients. The topology of α-synuclein pathology and its chronological change are known to be closely correlated with the symptomatic phenotypes^10–12^, indicating the local neurotoxicity of aggregated α-synuclein molecules. In the marmoset model of α-synuclein propagation, intensification and expansion of α-synuclein depositions visualized by ^18^F-C05-05 were in association with declines of the nigral dopaminergic neurons and their striatal terminals as assessed by PET imaging of dopamine transporters, in resemblance to the dopaminergic deficits in PD. This observation also implies that ^18^F-C05-05 could illuminate α-synuclein species critically involved in functional and structural disruptions of neurons. While our preliminary data in clinical PET assessments implied that progressive α-synuclein accumulations in the midbrain substantia nigra were traceable and could be associated with the advancement of motor deficits, it is yet to be examined whether ^18^F-C05-05-PET permits the pursuit of the α-synuclein dissemination along the clinical course of PD and DLB.

The total amount of abnormal α-synuclein proteins was reported to be approximately 50 – 200 nM in the brainstem and subcortical regions of advanced DLB and MSA cases, which was more than 10-fold smaller than the amount of Aβ peptide deposited in the brain of AD patients.^48^ This finding raises a concern regarding the visibility of α-synuclein pathologies by PET in a clinical setting. The IC50 value of ^18^F-C05-05 for the homologous blockade of its binding to α-synuclein aggregates in DLB tissue was 5 – 6 times lower than those of ^11^C-PBB3 and ^18^F-PM-PBB3 for the self-blockade of their binding to tau fibrils in AD tissue. Accordingly, PET with ^18^F-C05-05 would visualize α-synuclein depositions in DLB cases even if the brains of these patients possess radioligand binding sites with 5-fold lower density than AD brains. In line with this notion, we observed overt intensification of PET signals in the midbrains of moderate and severe LBDs, in contrast to the unremarkable enhancement of the radioligand retention in a PD case with the mildest manifestations. For the sensitive detection of early-stage Lewy pathologies, a compound with IC50 below 1 nM might be required and would be generated by further structural modifications of the C05 series chemical. An alternative strategy for gaining the detectability of nigral lesions may be the employment of a high-resolution PET scanner, as exemplified by a lately approved head-dedicated imaging device.^49^

In addition to the reactivity with the target lesion, the entry of the compound into the brain is a crucial factor for yielding a high signal-to-noise ratio in PET neuroimaging. Among C05 series compounds, C05-01 and C05-05 were fluorinated chemicals with desirable *in vitro* binding properties (Figure 1D).^33^ However, visualization of α-synuclein aggregates by intravital two-photon microscopic imaging of model mice was unsuccessful with C05-01. It is conceivable that the hydroxy moiety of C05-01 could be promptly conjugated with sulfate after systemic administration in a mode similar to PBB3^50^, and such metabolic modification was circumvented by the replacement of this structural group with a fluoroisopropanol moiety in PM-PBB3^19^ and C05-05 (Figure S11), increasing the amount of the intact compound entering the brain (compare Figures 3B and S12).

Another pharmacokinetic property affecting the *in vivo* target contrast is the rate of the radioligand washout from the brain. Radioligands with a prompt clearance, such as ^18^F-SPAL-T-06, yield low background signals, while a significant amount of free compounds could be eliminated from the brain tissue before a pseudo-equilibrium of the specific binding is reached. The relatively slow washout of ^18^F-C05-05 from the brain is supposedly advantageous for its efficient binding to α-synuclein deposits, enabling the detection of midbrain pathologies in LBDs. The retarded brain kinetics of ^18^F-C05-05 was elicited by its increased lipophilicity relative to ^18^F-SPAL-T-06, which also gave rise to augmented non-specific radioligand binding to myelin components. Indeed, radioligand accumulations in white matter regions were observed in ^18^F-C05-05-PET imaging of a marmoset model even before inoculation of α-synuclein fibrils. The clinical PET analysis of the substantia nigra was not profoundly influenced by this ligand property due to relatively low myelin contents in this area, but the high-contrast visualization of Lewy pathologies might be hampered in other regions with high white matter fractions. Although we also acknowledged elevated ^18^F-C05-05 retentions in several subcortical structures of MSA brains, the difference in SUVRs between the MSA and control subjects did not exceed the values in ^18^F-SPAL-T-06-PET, which conceivably stemmed from the abundant localization of GCIs in myelin-rich areas. In consideration of these issues, more sensitive α-synuclein imaging would be achieved by generating a compound with lipophilicity and clearance rate between these two PET probes. Moreover, several tau PET ligands show off-target binding to monoamine oxidases A and B.^51–53^ By contrast, previous studies documented that ^11^C-PBB3 and its analogs, including ^18^F-PM-PBB3 and ^18^F-C05-01, did not cross-react with these enzymes^19,33,54^, and our *in vitro* binding assay suggested insensitivity of ^18^F-C05-05 to monoamine oxidases (Figure S13).

In the present study, we granted the highest priority to the sensitive PET detection of α-synuclein pathologies with a high-affinity radioligand and succeeded in visualizing Lewy pathologies in the substantia nigra. In the brains of α-synucleinopathy patients, α-synuclein lesions are often co-localized with Aβ and tau aggregates, but patients with amyloid β-positive LBDs were not contained in this study. Depositions of tau in the substantia nigra of PD and DLB cases are uncommon, whereas tau pathologies at Braak stage III or above and Aβ pathologies are observed in more than 50% and 80% of α-synucleinopathy patients, respectively.^55^ Indeed, two out of five LBD patients with presented increased ^18^F-PM-PBB3 retentions in the temporal cortex (cases 2 and 5 in Figure S8). This finding raises the necessity for the development of a specific ligand for α-synuclein deposits with minimal cross-reactivity with other pathological fibrils. ^18^F-C05-05 displayed more than eight times smaller IC50 (1.5 nM) in DLB homogenates than in AD homogenates (12.9 nM), but its reactivity with tau deposits might not be markedly lower than that of ^11^C-PBB3 and ^18^F-PM-PBB3. In view of the putative structure-activity relationships indicated in this study, however, we are able to take advantage of β-sheet ligands with the (*E*)-hex-2-en-4-yne linker as potent binders, and structural modifications will be made for enhancing the selectivity of the chemicals by replacing aromatic rings and sidechains. It is also of significance that optical and PET imaging modalities can be utilized for the characterization of new candidate imaging agents. In addition, a recent non-clinical study has proposed a distinct chemical class with high selectivity for α-synuclein versus Aβ and tau fibrils, although the clinical applicability of the new probe remains to be determined.^56^ The substructures of these compounds would be implemented in the C05 series for reducing the cross-reactivity with non-α-synuclein components.

To conclude, the current neuroimaging platform incorporating C05-05 is implementable for multi-scale analysis of the neurodegenerative α-synuclein fibrillogenesis and pharmacological actions of a drug candidate on this etiological process in animal models. Our assays have also provided essential information on the feasibility of ^18^F-C05-05 as the first clinically available PET probe for Lewy-type α-synuclein pathologies in humans.

## ACKNOWLEDGEMENTS

The authors thank Asumi Orihara, Masaki Oya, Hideki Matsumoto, Shin Kurose, Masanori Ichihashi, Soichiro Kitamura, Sho Moriguchi, Takahiko Tokuda, Asaka Oyama, Hong Zhang, Yuki Momota, Takeharu Minamihisamatsu, Shoko Uchida, Kana Osawa, Takahiro Shimizu, Tomomi Kokufuta, Jun Kamei, Ryuji Yamaguchi, Yuichi Matsuda, Yoshio Sugii, Anzu Maruyama, Takashi Okauchi, and the staff of the Department of Advanced Nuclear Medicine Sciences at the National Institutes for Quantum Science and Technology for technical assistance. The authors acknowledge support for the recruitment of patients by Dr. Atsuhiko Sugiyama at the Chiba University Hospital, Dr. Taku Hatano at the Juntendo University Hospital, and Dr. Miwako Kawamura at the Koshigaya Municipal Hospital. We thank Shunsuke Koga and Dennis W. Dickson at the Mayo Clinic, and John Robinson, John Q. Trojanowski, and Virginia M.-Y. Lee at the University of Pennsylvania for case selection and kindly sharing postmortem brain tissues, and APRINOIA Therapeutics for kindly sharing precursor of ^18^F-PM-PBB3. This study was supported in part by MEXT KAKENHI Grant Number JP22K07529, and by AMED under Grant Number JP18dm0207018, JP22dm0207072, JP21dk027046, JP22dk0207063, and by the Collaborative Research Project (2021-201907) of the Brain Research Institute, Niigata University.

## AUTHOR CONTRIBUTIONS

Conceptualization, M.O., H.E., and M.Higuchi; Formal Analysis, M.O., H.E., M.T., M.F., Y.N., and K.Mimura; Investigation, M.O., H.E., Y.T., K.Matsuoka, M.T., K.Tagai, Y.Kataoka, K.H., K.Takahata, C.S., N.K., M.F., W.M., Y.N., K.Mimura, K.Kumata, T.K., S.K.M., Y.Y., H.Shimizu, and K.Kawamura; Resources, M.Hasegawa, A.S., H.T., H.Shimizu, and A.K.; Visualization, M.O., H.E., M.T., Y.N., and K.Mimura; Supervision, H.Shinotoh, H.Shimada, Y.Kimura, M.I., T.S., T.M., N.S., M.-R.Z., M.Hasegawa, and M.Higuchi; Project Administration, M.Higuchi; Funding Acquisition, M.O., and M.Higuchi; Critical Review and Editing of the Manuscript, all authors.

## STAR★METHODS

KEY RESOURCES TABLE.

STAR★METHODS: KEY RESOURCES TABLE
| REAGENT or RESOURCE | SOURCE | IDENTIFIER |
| --- | --- | --- |
| Antibodies |  |  |
| pS129 | Abcam | Cat.#:ab59264;<br>RRID:AB_2270761 |
| LB509 | Abcam | Cat.#:ab27766;<br>RRID:AB_727020 |
| AT8 | ThermoFisher Scientific | Cat.#:MN1020;<br>RRID:AB_223647 |
| 6E10 | BioLegend | Cat.#:803004;<br>RRID:AB_2715854 |
| pS199/202 | ThermoFisher Scientific | Cat.#:44-768G;<br>RRID:AB_2533749 |
| Chemicals, Peptides, and Recombinant Proteins |  |  |
| <sup>18</sup>F-C05-05 | This paper | N/A |
| <sup>11</sup>C-PBB3 | Maruyama et al., 2013 | N/A |
| <sup>18</sup>F-PM-PBB3 | Tagai et al., 2021 | N/A |
| <sup>11</sup>C-PE2I | Ando et al., 2012 | N/A |
| <sup>18</sup>F-C05-01 | Miranda-Azpiazu et al., 2020 | N/A |
| C05-03 | This paper | N/A |
| BF-227 | Nard Institute | Cat. #: NP039-0 |
| sulforhodamine 101 | Sigma-Aldrich | Cat. #: S7635 |
| Recombinant mouse wild-type α-synuclein and fibrils | Masuda-Suzukake et al., 2014 | N/A |
| Recombinant marmoset wild-type α-synuclein and fibrils | This paper | N/A |
| Experimental Models: Organisms/Strains |  |  |
| α-Syn mouse | Masuda-Suzukake et al., 2014 | N/A |
| α-Syn marmoset | this paper | N/A |
| Biological Samples |  |  |
| Table S1 | this paper | N/A |
| Software and Algorithms |  |  |
| Prism | Graph Pad | https://www.graphpad.com/scientific-software/prism/ |
| PMOD | PMOD Technologies | https://www.pmod.com/web/ |
| Statistical Parametric Mapping software | The Wellcome Centre for Human Neuroimaging | https://www.fil.ion.ucl.ac.uk/spm/ |
| MATLAB | MathWorks | https://www.mathworks.com/ |
| MVision brain | M Corporation | https://www.corporate-m.com/ |
| MRICron | Chris Rordens<br>Neuropsychology Lab | https://www.nitrc.org/projects/mricron |
| ImageJ (FIJI) | NIH | https://fiji.sc/ |

## RESOURCE AVAILABILITY

### Lead Contact

Further information and requests for resources and reagents should be directed to the Lead Contact, Hironobu Endo, Maiko Ono, and Makoto Higuchi.

## EXPERIMENTAL MODEL AND SUBJECT DETAILS

### Experimental animals

All animals studied here were maintained and handled in accordance with the National Research Council’s Guide for the Care and Use of Laboratory Animals. Protocols for the present animal experiments were approved by the Animal Ethics Committees of the National Institutes for Quantum Science and Technology (approval number: 07-1049-31, 11-1038-11). A total of 35 adult C57BL/6J mice (male, mean age 5.4 months, Japan SLC Inc) were used for the histochemical analysis, *ex vivo* examination, two-photon microscopy and PET scanning, and one adult marmoset (male, 2 years old, 300-365 g body weights) was used for PET scanning and histochemical analysis in this study. All mice and the marmoset were maintained in a 12 hours’ light/dark cycle with ad libitum access to standard diet and water.

### Human subjects

We included eight HCs, five patients with Lewy body diseases (PD and DLB spectrum) and three patients with MSA-P in the present study. All HCs (4 male, mean age 66.1 years) had no history of neurologic and psychiatric disorders. Four PD patients and one DLB patient (4 male, mean age 67.8 years) met UK Parkinson’s disease society brain bank clinical diagnostic criteria or McKeith’s criteria.^57,58^ Three patients with MSA (1 male, mean age 72.3 years) were clinically diagnosed according to the Gilman’s criteria.^59^ Two of the three MSA-P cases also showed mild cerebellar ataxia. The demographic profiles of HCs, PD/DLB and MSA patients are summarized in Table 1 and Table S2. When comparing both sexes between groups, there was a significant difference between HCs and PD/DLB (Table 1) and no significant difference between the three groups (Table S2). Individuals with PiB positivity and increased PM-PBB3 retentions following Braak tau stages were excluded from the present study as their brains were conceived to harbor AD pathologies. In this assessment, visual reads of PET images were conducted by three raters, H.E., K.Matsuoka and K.Tagai, according to the established criteria for PiB PET image^60^ and practical expertise with more than three years of reading experience in PM-PBB3-PET assays. In addition, we also excluded subjects with enhanced PM-PBB3 radiosignals in the subthalamic nucleus and neighboring subcortical areas and several cortical regions indicative of PSP and other frontotemporal lobar degeneration. Clinical diagnoses of patients were also validated according to their neurological examinations by H.E., K.H. and Y.Kataoka. Written informed consents were obtained from all subjects and/or from spouses or other close family members if subjects were cognitively impaired. This study was approved by the Radiation Drug Safety Committee and National Institutes for Quantum Science and Technology Certified Review Board of Japan. The study was registered with Japan registry of clinical trials (jRCTs031210180).

## METHOD DETAILS

### Compounds

**(*E*)-2-(4-(6-(methylamino)pyridin-3-yl)but-1-en-3-yn-1-yl)benzo[*d*]thiazol-6-ol** (C05-03) (^1^H nuclear magnetic resonance (NMR) (400 MHz, DMSO-D_6_): δ 9.97 (s, 1H), 8.19 (d, *J* = 4.0 Hz, 1H), 7.77 (d, *J* = 8.0 Hz, 1H), 7.48 (dd, *J* = 8.0, 4.0 Hz, 1H), 7.36 (d, *J* = 4.0 Hz, 1H), 7.19 (s, 1H), 7.15 (s, 1H), 7.13-7.10 (m, 1H), 6.97 (dd, *J* = 8.0, 4.0 Hz, 1H), 6.84 (d, *J* = 16 Hz, 1H), 6.48 (d, *J* = 8.0 Hz, 1H), 2.81 (d, *J* = 4.0 Hz, 3H)) and **(*E*)-1-fluoro-3-((2-(4-(6-(methylamino)pyridine-3-yl)but-1-en-3-yn-1-yl)benzo[*d*]thiazol-6-yl)oxy)propan-2-ol** (C05-05) (^1^H NMR (400 MHz, DMSO-D_6_): δ 8.19 (d, *J* = 2.3 Hz, 1H), 7.87 (d, *J* = 9.2 Hz, 1H), 7.69 (d, *J* = 2.7 Hz, 1H), 7.48 (dd, *J* = 8.7, 2.3 Hz, 1H), 7.20 (d, *J* = 16.0 Hz, 1H), 7.15-7.11 (m, 2H), 6.91 (d, *J* = 16.0 Hz, 1H), 6.48 (d, *J* = 8.2 Hz, 1H), 5.49 (d, *J* = 5.0 Hz, 1H), 4.60-4.53 (m, 1H), 4.50-4.40 (m, 1H), 4.12-4.00 (m, 3H), 2.81 (d, *J* = 4.6 Hz, 3H)) were custom-synthesized (Nard Institute). NMR spectra were obtained on a JEOL ECS-400 spectrometer at 400 MHz. Chemical shifts (δ) are reported in ppm, and coupling constants are reported in Hertz. The multiplicities are abbreviated as follows: s = singlet, d = doublet, m = multiplet, and dd = doublet of doublets (Figure S15). C05-01^33^, tosylate precursor of ^18^F-C05-05, PM-PBB3^19^, and tosylate precursor of ^18^F-PM-PBB3^19^ were custom-synthesized (Nard Institute and KNC Laboratories). PBB3 (Nard Institute, NP076-0), desmethyl precursor of ^11^C-PBB3 (Nard Institute, NP076-2), desmethyl precursor of ^11^C-PE2I (Nard Institute, PP022-1), BF-227 (Nard Institute, NP039-0), clorgiline (Sigma-Aldrich, M3778), selegiline (Sigma-Aldrich, NMID822), and sulforhodamine 101 (Sigma-Aldrich, S7635) are commercially available. All experiments with C05-01, C05-03, C05-05, ^18^F-C05-05, PBB3, ^11^C-PBB3, PM-PBB3, and ^18^F-PM-PBB3 were performed under UV-cut light to avoid photo-isomerization of these compounds.^19^

### Antibodies

Monoclonal antibodies against α-synuclein phosphorylated at Ser 129 (pS129; abcam, ab59264), α-synuclein (LB509; abcam, ab27766), tau phosphorylated at Ser 202 and Thr 205 (AT8; ThermoFisher Scientific, MN1020), and amyloid β (6E10; BioLegend, 803004), and polyclonal antibody against tau phosphorylated at Ser 199 and Thr 202 (pS199/202; ThermoFisher Scientific, 44-768G) are commercially available.

### Postmortem brain tissues

All brain autopsies were performed after consent of the legal next of kin or an individual with power of attorney. Postmortem human brains were obtained from autopsies carried out at the Department of Neuroscience of the Mayo Clinic on patients with DLB and MSA, at the Center for Neurodegenerative Disease Research of the University of Pennsylvania Perelman School of Medicine on patients with AD, and at the Department of Pathology of Niigata University on PDD, MSA-P, and control cases. Studies of autopsy samples were approved by the National Institutes for Quantum Science and Technology Certified Review Board (approval number: 14-015). Tissues for homogenate binding assays were frozen, and tissues for histochemical, immunohistochemical and autoradiographic labeling were frozen or fixed in 10% neutral buffered formalin followed by embedding in paraffin blocks.

### Preparation of recombinant α-synuclein and fibrils

Recombinant mouse and marmoset wild-type α-synuclein and fibrils were prepared as described previously.^8,37^ Briefly, purified α-synuclein (7 –10 mg/mL) was incubated at 37°C in a shaking incubator at 200 rpm in 30 mM Tris-HCl, pH 7.5, containing 0.1% NaN_3_, for 72 hours. α-synuclein fibrils were pelleted by spinning the assembly mixtures at 113,000 × g for 20 min, resuspended in saline, and sonicated for 3 min (Biomic 7040 Ultrasonic Processor, Seiko). Protein concentrations were determined by high performance liquid chromatography (HPLC) and adjusted to 4 mg/mL by dilution with saline.

### Stereotaxic surgery

For histochemistry, *ex vivo* examination and *in vivo* longitudinal imaging by two-photon laser scanning, nine-week-old mice anesthetized with 1.5% (v/v) isoflurane were unilaterally injected with 3 μL of recombinant mouse α-synuclein fibrils or 3 μL of saline into striatum (Interaural 3.82 mm, Lateral 2.0 mm, Depth 2.0 mm) via glass capillary.^8^ For PET study and *ex vivo* autoradiography, nine-week-old mice anesthetized with 1.5% (v/v) isoflurane were bilaterally injected with 3 μL of recombinant mouse α-synuclein fibrils or 3 μL of saline into striatum. For *in vivo* evaluation of ligands by two-photon laser scanning, mice anesthetized with 1.5% (v/v) isoflurane were unilaterally injected with 3 μL of recombinant mouse α-synuclein fibrils into somatosensory cortex (Interaural 1.98 mm, Lateral 2.5 mm, Depth 0.375 mm).

In the marmoset, surgeries were performed under aseptic conditions in fully equipped operating suite. We monitored body temperature, heart rate and SpO_2_ throughout all surgical procedures. The marmoset (2 years old at the time of surgery) was immobilized by intramuscular injection of ketamine (25 mg/kg) and xylazine (2 mg/kg) and then intubated by endotracheal tube. Anesthesia was maintained with isoflurane (1-3%, to effect). Prior to surgery, magnetic resonance (MR) imaging (20 cm bore, Biospec, Avance-III system; Bruker BioSpin) and X-ray computed tomography (CT) scans (Accuitomo170, J. MORITA CO.) were performed under anesthesia (isoflurane 1-3%, to effect). Overlay MR and CT images were created using PMOD image analysis software (PMOD Technologies Ltd) to estimate stereotaxic coordinates of target brain structures. For injections, the marmoset underwent surgical procedure to open burr holes (∼3 mm diameter) for the injection needle. Recombinant marmoset α-synuclein fibrils (right hemisphere, total 100 µl; 50 µl × 2 regions) and saline (left hemisphere, total 100 µl; 50 µl × 2 regions) were pressure-injected into caudate nucleus (Interaural 9.75 mm) and putamen (Interaural 9.75 mm) by Hamilton syringe mounted into motorized microinjector (UMP3T-2, WPI) held by manipulator (Model 1460, David Kopf, Ltd.) on a stereotaxic frame.^37^

### Two-photon laser-scanning microscopy

For surgical procedure, animals were anesthetized with a mixture of air, oxygen and isoflurane (3-5% W/V for induction and 2% W/V for surgery) via a facemask, and a cranial window (4.5-5 mm in diameter) was attached over the right somatosensory cortex, centered at 2.5 mm caudal and 2.5 mm lateral from bregma.^61^ Two-photon imaging was performed in awake mice two weeks after cranial window surgery at the earliest. Sulforhodamine 101 dissolved in saline (5 mM) was intraperitoneally administered (4 µl/g body weight) just before initiation of imaging experiments, and 0.05 mg of C05-05, C05-01, and PBB3 dissolved in dimethyl sulfoxide: saline = 1:1 (0.05% W/V) was intraperitoneally administered at various time points. Animals were placed on a custom-made apparatus, and real-time imaging was conducted by two-photon laser scanning microscopy (TCS-SP5 MP, Leica) with an excitation wavelength of 850-900 nm. In evaluation of *in vivo* labeling of α-synuclein inclusions with ligands, two-photon imaging was performed before and 5, 30, 60, and 90 min after administration of ligands. In *in vivo* longitudinal imaging of α-synuclein inclusions with C05-05, two-photon imaging was performed 90 min after administration of C05-05. An emission signal was separated by beam splitter (560/10 nm) and simultaneously detected through band-pass filter for ligands (525/50 nm) and sulforhodamine 101 (610/75 nm). A single image plane consisted of 1024×1024 pixels, with in-plane pixel size of 0.45 μm. Volume images were acquired up to maximum depth of 200-500 μm from cortical surface with z-step size of 2.5 μm.

### Radiosynthesis

^11^C-PE2I, ^11^C-PBB3, ^18^F-PM-PBB3, and ^18^F-C05-01 were radiosynthesized using their desmethyl, tosylate or nitro precursors as described previously.^16,19,62,63 18^F-C05-05 were radiosynthesized using their phenolic or tosylate precursors as described in Figure S4.^64^ Molar activity of ^18^F-C05-05 at the end of synthesis was 63-557 GBq/μmol, and ^18^F-C05-05 maintained its radioactive purity exceeding 90% for over 3 hours after formulation.

### PET imaging in mice and marmoset

PET scans were performed by microPET Focus 220 scanner (Siemens Medical Solutions). Mice were anesthetized with 1.5-2.0% isoflurane during all PET procedures. Emission scans were acquired for 90 and 120 min in 3D list mode with an energy window of 350-750 keV immediately after intravenous injection of ^18^F-C05-01 (23.5 ± 0.2 MBq) and ^18^F-C05-05 (30.8 ± 0.4 MBq), respectively. Images were reconstructed by either maximum a posteriori methods or filtered back projection (FBP) using a Hanning filter cut-off at Nyquist frequency (0.5 mm^-1^). All image data were subsequently analyzed using PMOD software (PMOD Technologies). For spatial alignment of PET images, template MR images generated previously^65^ were used in this study. Volumes of interest (VOIs) were manually placed on the striatum, cortex, amygdala and cerebellum. The marmoset was anesthetized with 1-3% isoflurane during all PET procedures. Transmission scans were performed for about 20 min with a Ge-68 source. Emission scans were acquired for 120 min and 90 min in 3D list mode with an energy window of 350-750 keV after intravenous bolus injection of ^18^F-C05-05 (89.6 ± 15.3 MBq) and ^11^C-PE2I (89.2 ± 2.0 MBq), respectively. PET images were reconstructed with FBP. All image data were subsequently analyzed using PMOD software. VOIs were placed on the caudate nucleus, putamen and cerebellum with reference to standard marmoset brain MR image.^66^

After anatomical standardization, decay-corrected time-activity curves in each of the VOIs were generated as the regional concentration of radioactivity averaged across the specific time window after radioligand injection. In the striatum and cortex, uptake value ratio to the cerebellum was calculated. To quantify ^11^C-PE2I binding, BP_ND_ was calculated with a simplified reference tissue model using the cerebellum as a reference region, and the caudate nucleus and the putamen as signal-rich regions.

### MRI studies in Human Subjects

MR images were acquired with a 3-T scanner, MAGNETOM Prisma (Siemens Healthcare). Three-dimensional volumetric acquisition of a T1-weighted gradient echo sequence produced a gapless series of thin sagittal sections (1-mm-thick sections, TE = 1.98 ms, TR = 2300 ms, TI = 900 ms, flip angle = 9°, field of view [FOV] = 250 mm, matrix size = 512 × 512 × 176).

### PET imaging in Human Subjects

PET scans were conducted with a Biograph mCT flow system (Siemens Healthcare; matrix size 400 × 400 × 109; voxel size [mm] 1 × 1 × 2 for ^18^F-PM-PBB3, ^18^F-C05-05 and ^11^C-PiB ligands, and a discovery MI (GE Medical Systems; matrix size 128 × 128 × 89; voxel size [mm] 2 × 2 × 2.8) for ^18^F-PM-PBB3 and ^11^C-PiB ligand. PET images were reconstructed using a filtered back-projection (FBP) algorithm with a Hanning filter (4.0 mm full width at half maximum) and ordered subset expectation maximization (OSEM) algorithm with time-of-flight (TOF), 3 iterations and 21 subsets. PET images reconstructed with the FBP algorithm were used for ROI analysis, and PET images reconstructed with the OSEM algorithm were used for figures.

^18^F-C05-05 PET scans were performed with two steps of scan protocol. In the first protocol, dynamic PET scans with arterial blood sampling were performed with 120 min after injection (frames: 6 × 10 s, 3 × 20 s, 6 × 1 min, 4 × 3 min, 20 × 5 min). During dynamic scanning, breaks were provided as needed, taking into consideration the subject’s stress, and we took a break from imaging at 50-70 min in one healthy case. In the second protocol, a 30-min PET acquisition was performed 90-min after injections (6 × 5 min frames). ^18^F-C05-05 had an average injected dose of 177.7 ± 11.5 MBq with molar activity at the time of injection of 181.3 ± 82.4 GBq/μmol.

^11^C-PiB PET scans were performed following a previously reported protocol.^16,67,68^ ^11^C-PiB (injected dose: 531.9 ± 44.2 MBq, molar activity: 91.0 ± 50.7 GBq/μmol) PET scan was conducted with a 20-min acquisition 50 min after injections. Each participant received 20-min acquisition 90 min after intravenous injection of ^18^F-PM-PBB3 PET scan (injected dose: 182.3 ± 4.5 MBq, molar activity: 242.8 ± 113.0 GBq/μmol, 4 × 5 min frames). Because of secondary use of data from another study (UMIN000030248), there was a time lag of up to 3.5 years for ^18^F-PM-PBB3 PET and 3.4 years for ^11^C-PiB PET in the HC group, and up to 1.6 years for ^18^F-PM-PBB3 PET and 1.7 years for ^11^C-PiB PET in the LBDs group from the ^18^F-C05-05 PET scan.

To determine the unmetabolized ^18^F-C05-05 fraction in arterial plasma, radiometabolite analysis was performed. Arterial blood samples (2 mL each) taken at 3, 10, 20, 40, 60, 90, and 120 min were subjected to metabolite analysis by high performance liquid radiochromatography. Blood samples were placed on ice immediately after sampling to avoid further metabolism and centrifuged at 13,000 g at 4 °C for 2 min. An aliquot of 0.3 mL plasma was mixed with 1 mL acetonitrile/methanol = 2/1 (v/v), and centrifuged at 13,000 rpm at 4 °C for 2 min for deproteinization. Then mean radioactivity recovery was 88% (83-91%) in the first volunteer. Then an aliquot of the supernatant was injected into a reverse-phase HPLC system (JASCO Corporation). YMC-Triart C18 ExRS (5 μm, 10 mm i.d. × 10 mm, YMC) was used for a guard column and YMC-Triart C18 ExRS (5 μm, 10 mm i.d. × 150 mm, YMC) was used for an analytical column. The mobile phase was 90% acetonitrile/ 0.1 μmol/L ammonium acetate solution (47.5/52.5, v/v) at an isocratic condition, and the flow rate was 5.0 mL/min. Effluent radioactivity was detected with a home-made NaI (TI) scintillation detector system.^69^ The retention time of the radiochromatography peak of ^18^F-C05-05 was identified by the optical absorption of standard C05-05 at a detection wavelength of 390 nm. The unmetabolized fraction was calculated as the peak area ratio of unmetabolized ^18^F-C05-05 to the total peaks detected.

### Image analyses in Human Subjects

#### Data preprocessing

Data preprocessing was performed using PMOD 4.3 (PMOD Technologies LLC, Switzerland), Statistical Parametric Mapping software (SPM12, Wellcome Department of Cognitive Neurology) and MVision brain (M Corporation, Tokyo, Japan). To obtain the T1-weighted MR segmentation map, MVision brain was performed to down sample the matrix of T1WI from 512 × 512 × 176 to 256 × 256 × 176. Regarding VOI analyses, the whole brain was automatically parcellated into 287 regions using the MVision brain, a standalone version of the MR brain image analysis system: BrainGPS. The BrainGPS is an MRI analysis cloud platform developed based on a technique, known as the MRIcloud (https://mricloud.org/). We used ROIs for the parietal cortex, putamen, Substantia nigra, Cerebellar gray matter, deep white matter [combined superior corona radiata (SCR) and posterior corona radiata (PCR)] based on the 287 VOIs obtained from the Mvision brain (Figure S16). Partial volume correction was not performed in the present study. SUVR images were generated from averaged PET images at the following intervals: 50-70 min (^11^C-PiB), 90-110 min (^18^F-PM-PBB3) and 100-120 min (^18^F-C05-05) post injection, respectively. For SUVR ROI analysis, acquired PET images were rigidly coregistered to individual T1-weighted MR images analyzed by the Mvision brain. On the other hand, for TAC data plotting, the ROIs obtained from Mvision brain were inverted to PET space for analysis. We decided the target region was the substantial nigra based on pathological hypothesis.^70,71^ The Cerebellar gray matter was used as reference region for ^11^C-PiB as previously described.^19^ Since α-synuclein pathology is also known in cerebellum in LBDs^41,42^, the deep white matter (SCR and PCR) was chosen as a reference region that was free of neuronal cell bodies and could be widely obtained for α-synuclein evaluation using ^18^F-C05-05. Equilibrium was reached from 100 minutes after injection by visual assessment based on the time activity curve (Figure S6 C). For qualitative image evaluation of ^18^F-C05-05 and ^18^F-PM-PBB3, SUVR images were also generated for ^18^F-PM-PBB3 using the deep white matter as the reference region (SUVRdwm).

Regarding the average image creation for representative groups of 11C-PiB SUVRcerebellar gray matter, ^18^F-PM-PBB3 SUVRdwm and ^18^F-C05-05 SUVRdwm for HCs, LBDs and MSAs, each image (T1WI and SUVR) was spatially normalized to MNI (Montreal Neurologic Institute) space (using a brain template consisting of 6 probability maps of SPM12 at 1.5mm resolution, then additionally transformed using diffeomorphic deformation [ANTS: advanced normalization tools] and fitted to the T1 MR template) using PMOD. The normalized images were then smoothed with a Gaussian kernel at 5 mm full width at half maximum.

### Characteristics of ^18^F-C05-05 in Human Subjects

We explored uptake into the brain, and the dynamic range and distribution of specific binding of ^18^F-C05-05. A head-to-head visual comparison of ^18^F-C05-05, ^18^F-PM-PBB3 and ^11^C-PiB was conducted in the same individuals (Figure S8 for LBDs, and Figure S9 for HCs and MSA-P).

### Assessing α-synuclein Deposits Associated with LBDs

Group comparisons using ROI in substantial nigra were conducted; in addition, correlation analyses between SUVRdwm of the ROI and MDS-UPDRS (Movement Disorder Society revised Unified Parkinson’s Disease Rating Scale) part III scores were also performed in LBDs.

### Histological examination

Mice were deeply anesthetized and sacrificed by saline perfusion, and brains were subsequently dissected and fixed in 4% paraformaldehyde in phosphate buffered saline (PBS) overnight at 4°C. After cryo-protection in PBS containing 20% sucrose, brains were embedded and frozen in OCT compound (SaKuRa), and 20-μm thick fixed frozen sections were prepared by cryostat. The marmoset was deeply anesthetized with an overdose of sodium pentobarbital (80 mg/kg, intravenous administration) and transcardially perfused with saline at 4°C, followed by 4% paraformaldehyde in PBS, pH 7.4. The brain was removed from the skull, postfixed in the same fresh fixative overnight, saturated with 30% sucrose in phosphate buffer at 4°C, and then cut serially into 40-μm-thick sections on a freezing microtome. For fluorescence labeling with ligands, mouse fixed frozen sections, marmoset fixed sections, 6-μm thick deparaffinized postmortem human brain sections and 20-μm thick fresh frozen human brain sections post-fixed in 4% paraformaldehyde in PBS were incubated in 20% and 50% ethanol containing 30 μM ligand, respectively, at room temperature for 30 min. The samples were rinsed with 20% or 50% ethanol for 5 min, dipped into distilled water twice for 3 min each, and mounted in non-fluorescent mounting media (VECTASHIELD; Vector Laboratories). Fluorescence images were captured by a DM4000 microscope (Leica) equipped with custom filter cube (excitation band-pass at 414/23 nm and suppression low-pass with 458 nm cut-off) and BZ-X710 fluorescence microscope (KEYENCE) equipped with Filter set ET-ECFP (Chroma Technology). Sections labeled with ligands in fluorescence microscopy, *ex vivo* examination and autoradiography and their adjacent sections were immunostained with pS129, LB509, pS199/202, and AT8 antibodies with antigen retrieval by autoclaving, and with 6E10 antibodies with antigen retrieval by formic acid. Immunolabeling was then examined using DM4000 and BZ-X710. Images were analyzed using ImageJ software (NIH Image).

### Autoradiography

*In vitro* autoradiography was performed using 6-µm-thick deparaffinized sections derived from MSA brains, 20-µm-thick fresh frozen sections post-fixed in 4% paraformaldehyde in PBS derived from DLB, PDD, MSA-P, and control brains, and 40-μm-thick fixed sections derived from α-Syn marmoset. For labeling with ^18^F-C05-05, sections were pre-incubated in 50 mM Tris-HCl buffer, pH 7.4, containing 20% ethanol at room temperature for 30 min, and incubated in 50 mM Tris-HCl buffer, pH 7.4, containing 20% ethanol and ^18^F-C05-05 (5 nM; molar activity: 557 GBq/µmol, 10 nM; molar activity: 260 GBq/µmol) at room temperature for 60 min. Excess concentration (10 µM) of C05-05 was added to the reaction to determine non-specific radioligand binding. The samples were then rinsed with ice-cold Tris-HCl buffer containing 20% ethanol twice for 2 min, and dipped into ice-cold water for 10 sec. The sections labeled with ^18^F-C05-05 were subsequently dried with warm air, and exposed to an imaging plate (BAS-MS2025; Fuji Film). *Ex vivo* autoradiography was performed in mice used in PET imaging. Mice were anesthetized with 1.5% (v/v) isoflurane and intravenously administered ^18^F-C05-05 (27.8 ± 0.2 MBq, molar activity: 63 GBq/µmol). 90 min after administration of ^18^F-C05-05, mice were then sacrificed by cervical dislocation, and the brains were removed. After quick freezing with powdered dry ice, 20-μm thick frozen sections were prepared by cryostat and exposed to an imaging plate. The imaging plate was scanned by BAS-5000 system (Fuji Film) to acquire autoradiograms. Images were analyzed using Multi Gauge software (Fuji Film).

### *In vitro* binding assay

Frozen tissues derived from the amygdala of a DLB patient, frontal cortex of an AD patient, and putamen of an MSA-P patient were homogenized in 50 mM Tris-HCl buffer, pH 7.4, containing protease inhibitor cocktail (cOmplete^TM^, EDTA-free; Roche), and stored at –80°C until analyses. To assay radioligand binding with homologous blockade, these homogenates (100 µg tissue) were incubated with 1 or 5 nM ^18^F-C05-05 (1 nM; molar activity: 557 GBq/µmol, 5 nM; molar activity: 63 GBq/µmol), 5 nM ^11^C-PBB3 (molar activity: 93.4 ± 24 GBq/µmol), or 1 nM ^18^F-PM-PBB3 (molar activity: 183.3 ± 89.2 GBq/µmol) in the absence or presence of non-radiolabeled C05-05, PBB3, or PM-PBB3 at varying concentrations ranging from 1×10^-11^ to 5×10^-7^ M in Tris-HCl buffer containing 10% ethanol, pH 7.4, for 30 min at room temperature. Non-specific binding of ^18^F-C05-05, ^11^C-PBB3, and ^18^F-PM-PBB3 was determined in the presence of 1×10^-7^ or 5×10^-7^ M C05-05, PBB3, and PM-PBB3, respectively. Samples were run in triplicate or quadruplicate. To assay radioligand binding with heterologous blockade, the DLB amygdala and AD frontal cortex homogenates (100 µg tissue) were incubated with 5 nM ^18^F-C05-05 (molar activity: 201 GBq/µmol) in the absence or presence of clorgiline and selegiline at varying concentrations ranging from 1×10^-11^ to 5×10^-7^ M in Tris-HCl buffer containing 10% ethanol, pH 7.4, for 30 min at room temperature. Concentration of the competitor inducing 50% inhibition (IC50) was determined by using non-linear regression to fit a concentration-binding plot to one-site and two-site binding models derived from the Cheng-Prusoff equation with GraphPad Prism version 5.0 (GraphPad Software), followed by F-test for model selection.

### *Ex vivo* fluorescence examination

Mice were anesthetized with 1.5% (v/v) isoflurane and intraperitoneally administered C05-05 (1.66 mg/kg). 90 min after administration of C05-05, mice were then sacrificed by cervical dislocation, and brains were removed. After quick freezing by powdered dry ice, 20-μm thick frozen sections were prepared by cryostat and mounted in VECTASHIELD. Fluorescence images of brain section with no additional staining were captured by DM4000 microscope equipped with custom filter cube and BZ-X710 fluorescence microscope equipped with Filter set ET-ECFP. For immunostaining, sections used for *ex vivo* examination were fixed in 4% paraformaldehyde in PBS overnight at 4°C just prior to staining.

### Molecular docking

Structure of type I (PDB ID: 6XYO) and Type II (PDB IDs: 6XYP and 6XYQ) filaments of α-synuclein from MSA was taken from Protein Data Bank.^6^ Hydrogens were added and energy minimized by protein preparation wizard of Maestro. Ligand structures were modeled using program LigPrep. Ligands were first docked blindly to five type I α-synuclein filaments by AutoDock Vina.^72^ A total of 200 docking conformations were generated and analyzed. From this blind docking, 14 binding regions were identified and then ligands were docked into each identified binding site individually. Docking in each binding site was performed by standard precision docking protocol of program Glide. Highest scoring binding pose of each ligand was analyzed and have been reported. Since site S10 and S11 are very close to each other, docking in these two was performed together in single docking run and labeled as S10-11. Binding site S8 and S12 were identified as most likely binding sites and were further analyzed in both the structures of type II filament. Docking scores were converted to affinities (*K*_d_) using relation Score = –*RT* ln (*K_d_*). S12 and S8 were compared by superimposing structures in PyMol.

### Radiometabolite analysis in mice

^18^F-C05-05 (23.2 ± 6.2 MBq, molar activity: 320.5 ± 10.6 GBq/µmol) was intravenously administered to awake C57BL/6J mice (male, six-month-old). Mice were decapitated at 5, 30, 60, and 90 min after ^18^F-C05-05 administration, and blood and brain samples were collected. For blood samples, the whole blood was centrifuged at 15,000×g for 2 min at 4°C. The resulting plasma (100 µL) was mixed with a 150 µL of acetonitrile/Methanol (100/50) solution and centrifuged at 15,000×g for 2 min at 4°C for deproteinization. An aliquot of the supernatant was then collected for HPLC analysis. For brain samples, the whole brain was homogenized in a 1 mL of ice-cold saline. The resulting homogenate (100 µL) was mixed with a 150 µL of acetonitrile/Methanol (100/50) solution and centrifuged at 15,000×g for 2 min at 4°C for deproteinization. The supernatant of each brain homogenate was subjected to HPLC analysis. The radiometabolites in the plasma and brain samples were analyzed by an HPLC system (JASCO Corporation) equipped with a high-sensitive positron detector (Ohyo Koken Kogyo). The column used was CAPCELL PAK C18 UG80 (5 µm, 4.6 mm i.d. × 250 mm, OSAKA Soda). The mobile phase was acetonitrile/50 mM ammonium acetate solution (55/45) at an isocratic condition, and the flow rate was 1.0 mL/min. The retention time of radioactivity corresponding to that of standard C05-05 detected at 390 nm was identified to be of ^18^F-C05-05. The fractions (%) of peak area corresponding to ^18^F-C05-05 and radiometabolites were calculated with decay correction.

## QUANTIFICATION AND STATISTICAL ANALYSIS

Statistical significance of the data was analyzed with GraphPad Prism version 5.0 or 8.0, and SPSS version 29. For comparison of two groups, we performed t-tests for items of continuous variables such as age and disease duration, and Man-Whitney tests for scores such as clinical data. For categorical data such as sex, we used the chi-squared test. For comparison of multiple groups, data were analyzed by one-way ANOVA with post-hoc Tukey’s HSD test for continuous variables and Kruskal-Wallis test with post-hoc Dunn’s test for clinical scores that were not normally distributed. For comparison of multiple groups and regions, data were analyzed by two-way repeated-measures ANOVA with Bonferroni’s post hoc analysis. Group comparison of SUVR values derived from region of interest (ROI) of substantial nigra between HCs and LBDs groups were conducted by two-sample t-test. To examine correlations between ^18^F-C05-05 SUVR in substantial nigra and MDS-UPDRS part III scores, we used Spearman’s rho test. Data in figures represent mean ± SD, with sample size in figure legends and detailed statistic values listed in Results.

## DATA AND SOFTWARE AVAILABILITY

Requests for data that support the finding of this study should be directed to the Lead Contact, Hironobu Endo, Maiko Ono, and Makoto Higuchi, and will be available upon reasonable request.

## SUPPLEMENTAL INFORMATION

Table S1. Antemortem clinical diagnosis and neuropathologic description of postmortem human brain tissues, Related to Figure 1, 5, 6.

Table S2. Demographics of individuals with HCs, LBDs and MSAs included in the present study, Related to Table 1.

Figure S1. Immunostaining of DLB and AD brain sections used for characterization of ligands, Related to Figure 1.

Figure S2. Temporal changes in the distribution of phosphorylated α-synuclein inclusions in mice injected with α-synuclein fibrils into the striatum, Related to Figure 2.

Figure S3. *Ex vivo* examination of brain sections from a mouse injected with α-synuclein fibrils and a control mouse, Related to Figure 2.

Figure S4. Radiosynthesis of ^18^F-C05-05, Related to Figure 3-7.

Figure S5. Binding mode of C05-05 in α-synuclein filaments predicted by molecular docking, Related to Figure 5.

Figure S6. PET data in representative cases, Related to Figure 6.

Figure S7. ^18^F-C05-05 PET images using ranges from 0, Related to Figure 6.

Figure S8. ^11^C-PiB and ^18^F-PM-PBB3 images of the same subject with ^18^F-C05-05 PET in patients with Lewy body disease (LBDs), Related to Figure 6.

Figure S9. ^11^C-PiB and ^18^F-PM-PBB3 images of the same subject with ^18^F-C05-05 PET in healthy controls (HCs) and patients with multiple system atrophy, parkinsonian (MSA-P), Related to Figure 6, 7.

Figure S10. Autoradiographic labeling of putamen sections derived from an MSA-P with ^18^F-C05-05, Related to Figure 7.

Figure S11. Metabolite analysis of ^18^F-C05-05 in mice, Related to Figure 3.

Figure S12. Time-radioactivity curve in the brain of wild-type mice in ^18^F-C05-01 PET, Related to Figure 3.

Figure S13. Inhibition of total binding of ^18^F-C05-05 by monoamine oxidase inhibitors in DLB and AD homogenates, Related to Figure 5.

Figure S14. Time-radioactivity curves in the brain of an α-Syn marmoset in ^18^F-C05-05 PET, Related to Figure 4.

Figure S15. ^1^H NMR spectra for C05-03 and C05-05, Related to Figure 1.

Figure S16. Representative reference and target region for PET standardized uptake value ratio (SUVR), Related to Figure 6.

