## Supplemental File for "Imaging α-synuclein pathologies in animal models and patients with Parkinson’s and related diseases"

**SUPPLEMENTAL INFORMATION**

**SUPPLEMENTAL FIGURES**

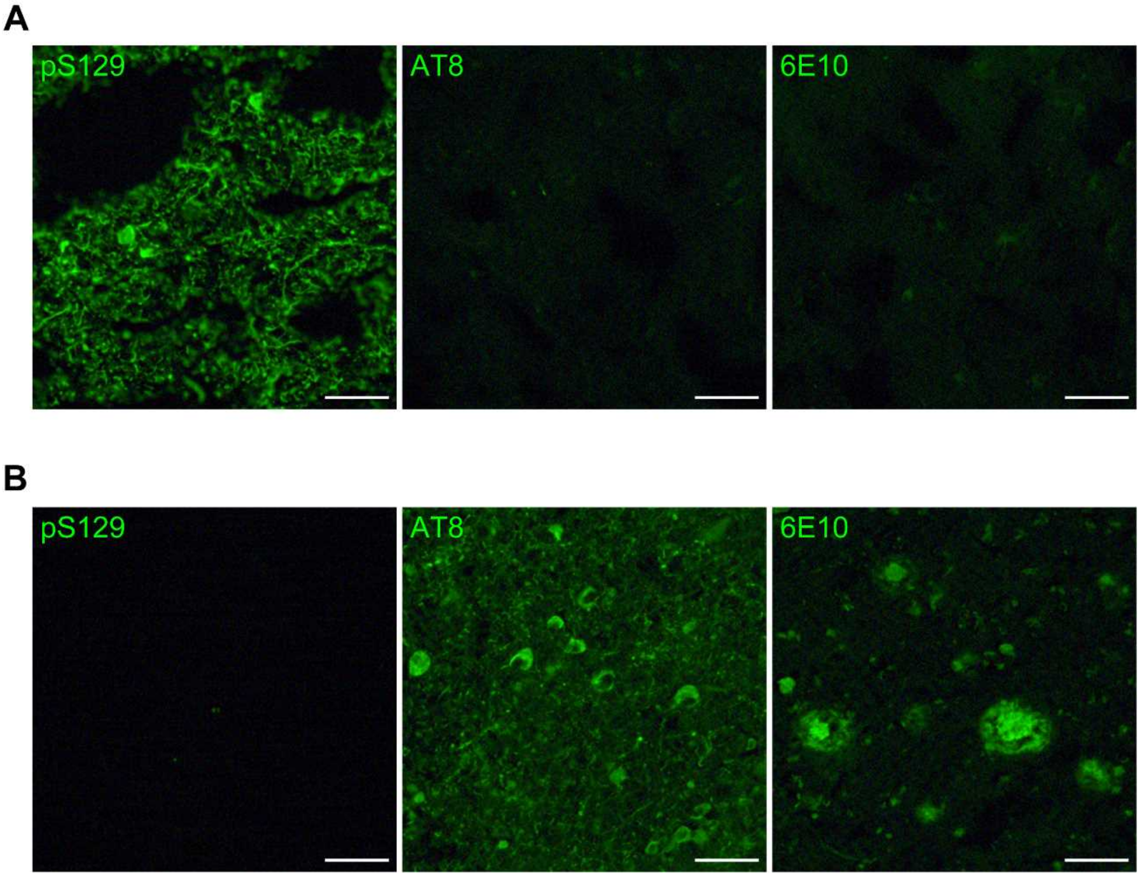


**Figure S1.** Immunostaining of DLB and AD brain sections used for characterization of ligands, Related to Figure 1.

(**A**) Immunostaining of amygdala sections derived from a DLB patient with pS129 (left), AT8 (middle), and 6E10 (right) confirms the absence of tau and Aβ deposit in this brain tissue. (**B**) Immunostaining of middle frontal gyrus sections derived from an AD patient with pS129 (left), AT8 (middle), and 6E10 (right) confirms the absence of α-synuclein deposits in this brain tissue.

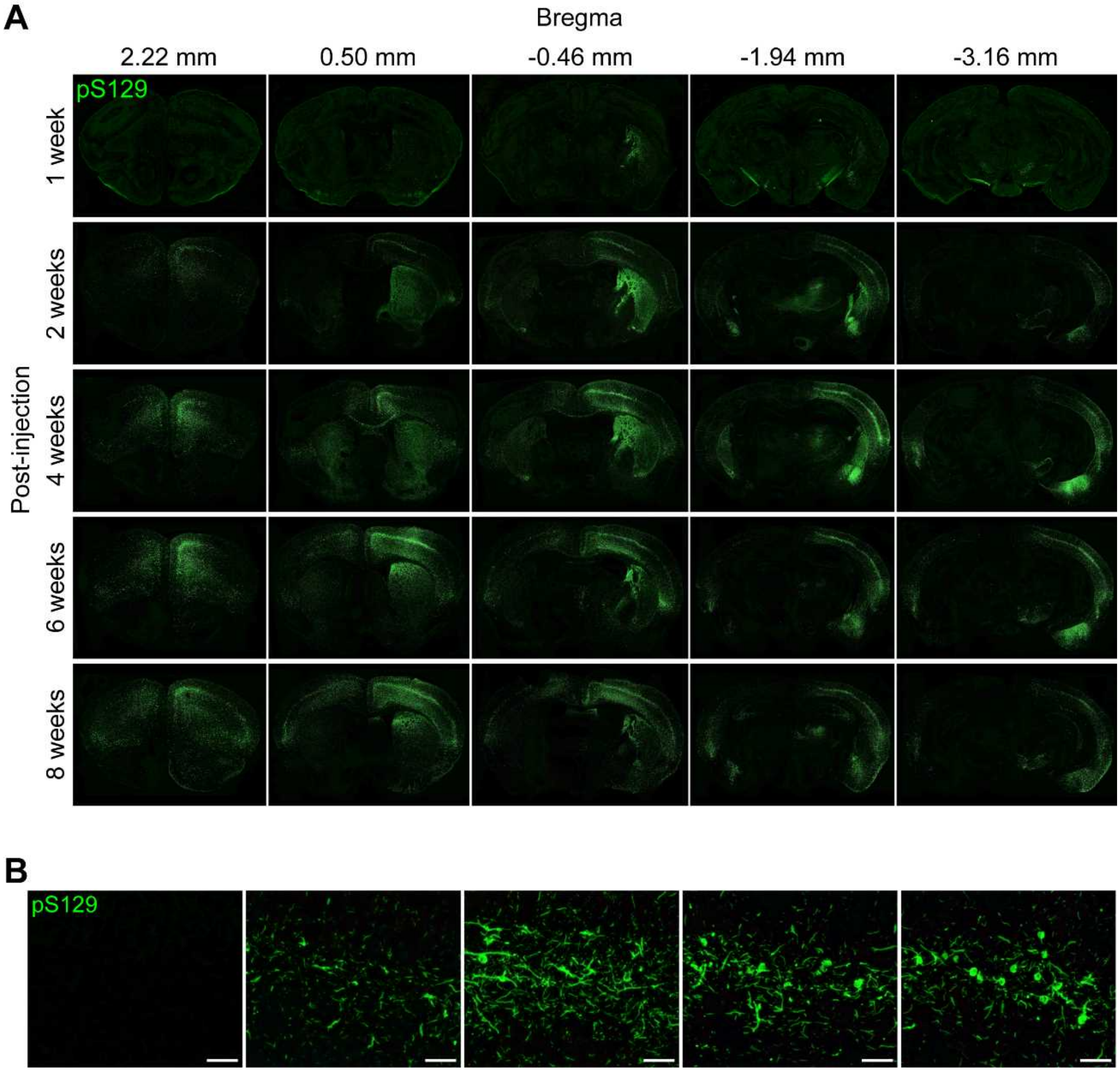


**Figure S2.** Temporal changes in the distribution of phosphorylated α-synuclein inclusions in mice injected with α-synuclein fibrils into the striatum, Related to Figure 2.

(**A, B**) Phosphorylated α-synuclein proteins were immunostained with pS129 in coronal brain sections at bregma +2.22, +0.5, -0.46, -1.94 and -3.16 mm derived from mice at 1, 2, 4, 6, and 8 weeks after inoculation of α-synuclein into the right striatum (**A**), and high-power photomicrographs of the ipsilateral somatosensory cortex (**B**). Scale bars, 50 μm.

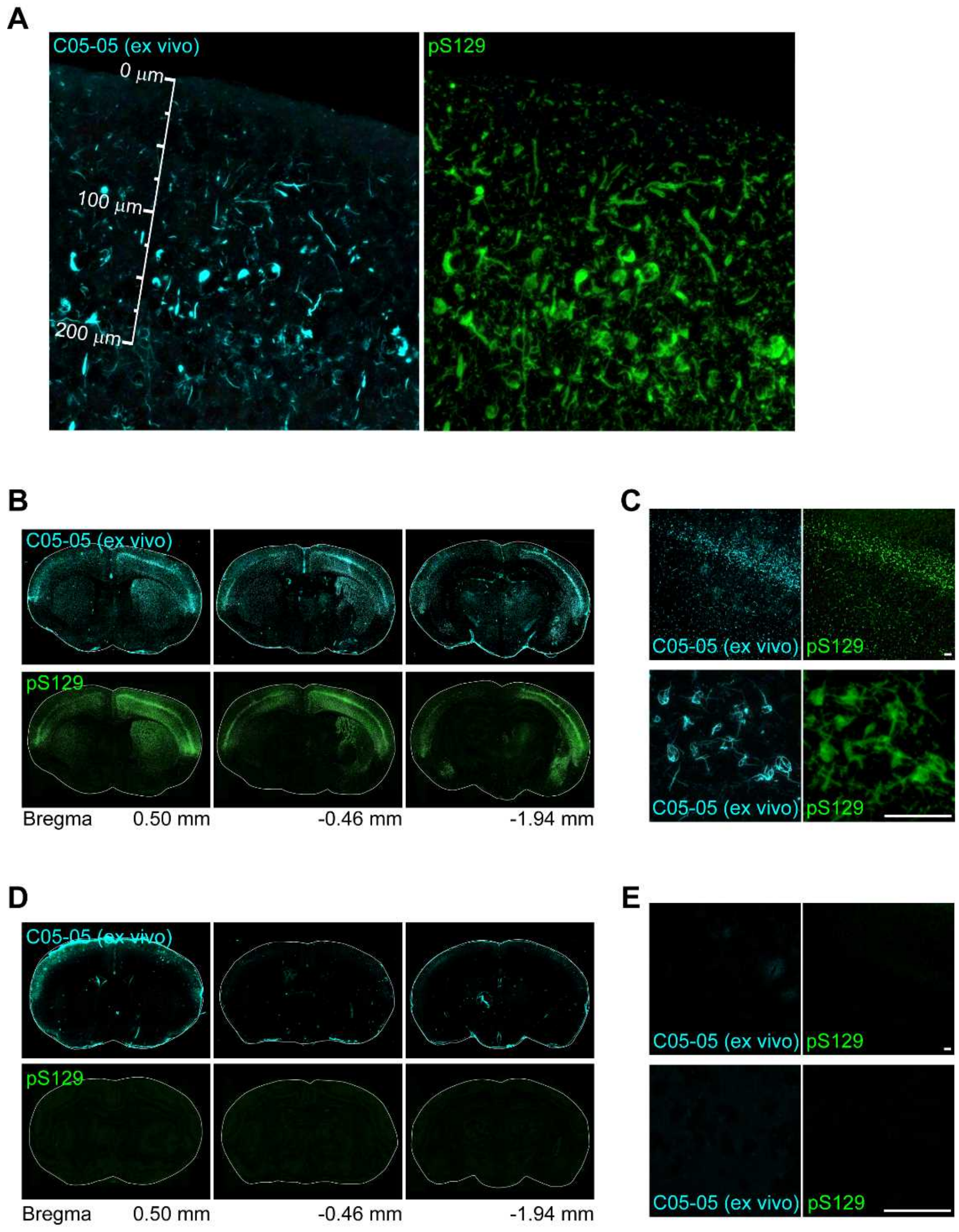


**Figure S3.** *Ex vivo* examination of brain sections from a mouse injected with α-synuclein fibrils and a control mouse, Related to Figure 2.

(**A**) *Ex vivo* examination of frozen brain sections from mouse injected with α-synuclein fibrils into cortex at 11 weeks after injection to detect intraperitoneally administered C05-05 (1.66 mg/kg) (left), and immunolabeling of adjacent brain section with pS129 (right). This α-Syn mouse was the same as used for *in vivo* two-photon laser microscopic imaging (Fig. 2). High-power photomicrographs of cortical sections demonstrated abundant accumulation of C05-05 positive-α-synuclein inclusions in somatosensory cortex of this α-Syn mouse. (**B**) *Ex vivo* examination of frozen brain sections from an α-Syn mouse at 8 weeks after the inoculation of α-synuclein fibrils into the right striatum. The brain tissue was collected at 90 min after intraperitoneal administration of C05-05 (1.66 mg/kg). Distributions of systemically injected C05-05 in coronal brain sections (top) and immunolabeling of adjacent brain sections with pS129 (bottom) at bregma +0.50, -0.46, and -1.94 mm are displayed. Brain samples collected from the α-Syn mouse at 90 min after intraperitoneal C05-05 injection contained numerous fibrillary inclusions labeled with the injected compound in broad areas of the brain, including the striatum, neocortex and amygdala, and these aggregates were subsequently stained with pS129. (**C**) Medium-power (top) and high-power (bottom) photomicrographs of cortical sections shown in **B**. Individual α-synuclein inclusions were found to be intensely labeled with intraperitoneally administered C05-05. (**D**) *Ex vivo* examination of frozen brain sections from a mouse at 8 weeks after injection of saline into the right striatum. The brain was taken at 90 min after intraperitoneal administration of C05-05 (1.66 mg/kg), and C05-05-derived signals (top) and immunolabeling with pS129 (bottom) were examined in adjacent brain sections. From the left, coronal brain sections at bregma +0.50, -0.46, and -1.94 mm are displayed. (**E**) Medium-power (top) and high-power (bottom) photomicrographs of cortical sections shown in **D**. *Ex vivo* examination revealed no apparent signals originating from of intraperitoneally administered C05-05 in the cerebral parenchyma of the control mouse. Scale bars, 50 μm (**C** and **E**).

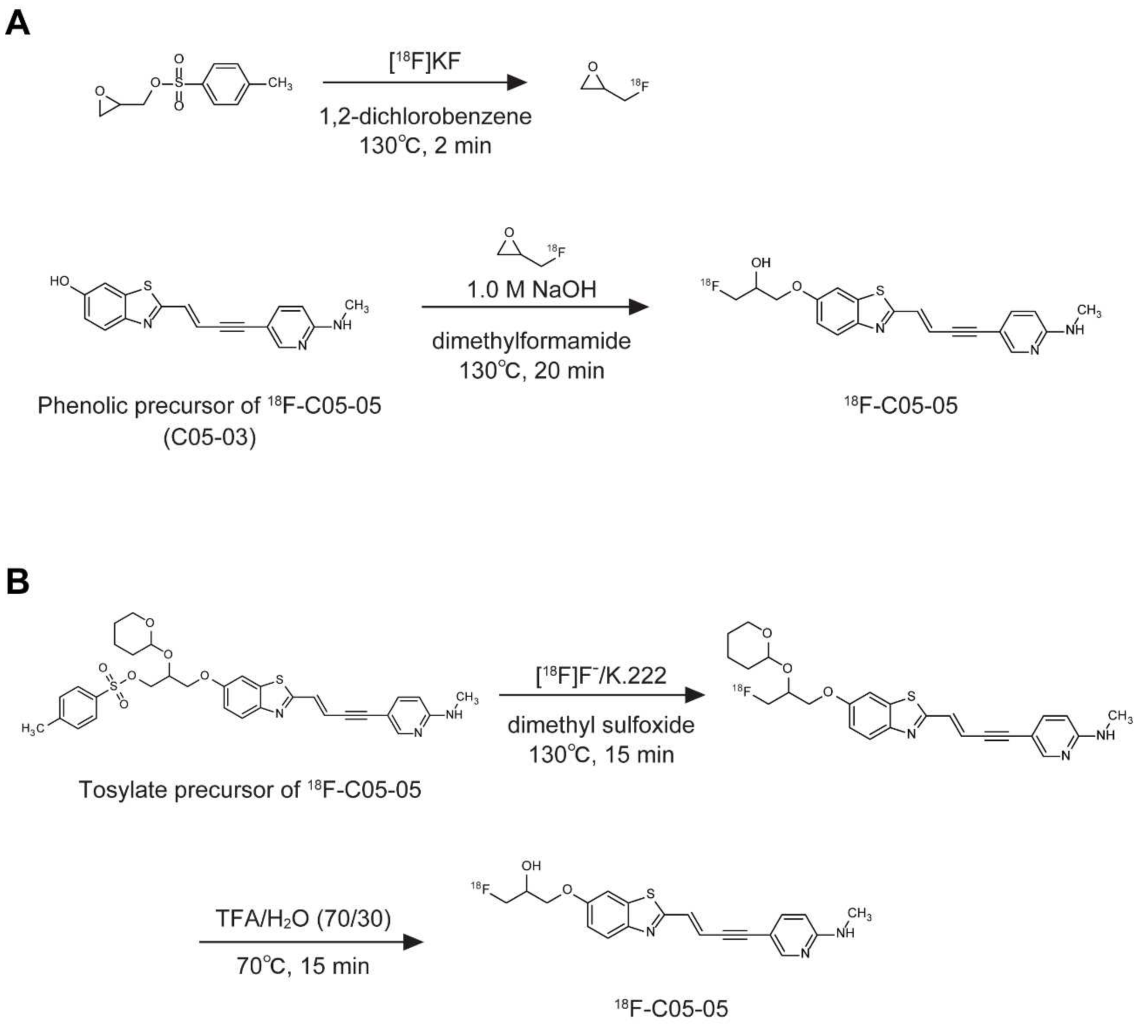


**Figure S4.** Radiosynthesis of $^{18}$F-C05-05, Related to Figure 3-7.

(**A**) Radiosynthesis of $^{18}$F-C05-05 using phenolic precursor (C05-03). Radiolabeling of $^{18}$F-C05-05 was accomplished by ring-opening reaction of (rac)-$^{18}$F-epifluorohydrin with a phenolic precursor in the presence of 1.0 M NaOH and dimethylformamide at 130°C for 20 min. After fluoroalkylation, the crude reaction mixture was transferred into a reservoir for preparative high performance liquid chromatography (HPLC) using Atlantis Prep T3 column (10 × 150 mm, Waters) with a mobile phase consisting of acetonitrile/water (25/75) with 0.1% trifluoroacetic acid (v/v) at a flow rate of 5 mL/min. The fraction corresponding to $^{18}$F-C05-05 was collected in a flask containing 100 μL of 25% ascorbic acid solution and Tween 80, and was evaporated to dryness under a vacuum. The residue was dissolved in 3 mL of saline (pH 7.4) to obtain $^{18}$F-C05-05. The final formulated product was chemically and radiochemically pure (>95%) as detected by analytical HPLC using Atlantis Prep T3 column (4.6 × 150 mm, Waters) with mobile phase consisting of acetonitrile/water (30/70) with 0.1% trifluoroacetic acid (v/v) at a flow rate of 1 mL/min. (**B**) Radiosynthesis of $^{18}$F-C05-05 using tosylated precursor. Tosylated precursor for $^{18}$F-labeling of C05-05 in dimethyl sulfoxide (DMSO) was reacted with [$^{18}$F]$^{18}$F$^{-}$ in the presence of $K_2CO_3$ and kryptfix 222 at 130°C for 15 min. After cooling the reaction vessel to 70°C, 70% trifluoroacetic acid solution was added to the mixture and maintained for 15 min to delete the protecting groups. The mixture of 4 mol/L sodium acetate solution and DMSO was added to the reaction vessel, and the mixture was transferred into a injector for semi-preparative HPLC (Semi-preparative HPLC condition: GL Sciences Inertsil ODS-HL column, 10 × 150 mm; acetonitrile/50 mM ammonium acetate = 50/50 as the mobile phase, 5 mL/min flow rate). The fraction corresponding to $^{18}$F-C05-05 was collected into a flask containing 25% ascorbic acid solution and Tween 80, and was evaporated to dryness under a vacuum. The residue was dissolved in saline and filtered with sterilization filter to obtain $^{18}$F-C05-05 as an injectable solution. The final formulated product was radiochemically pure (>95%) as detected by analytical radio-HPLC (analytical HPLC condition: ACE Excel 3 C18-Amide column, 4.6 × 100 mm; 90% aqueous acetonitrile mixed with 100 mmol/L ammonium phosphate buffer (pH 2.0) containing 5 mmol/L sodium 1-octanesulfonate = 35/65 as the mobile phase, 1 mL/min flow rate).

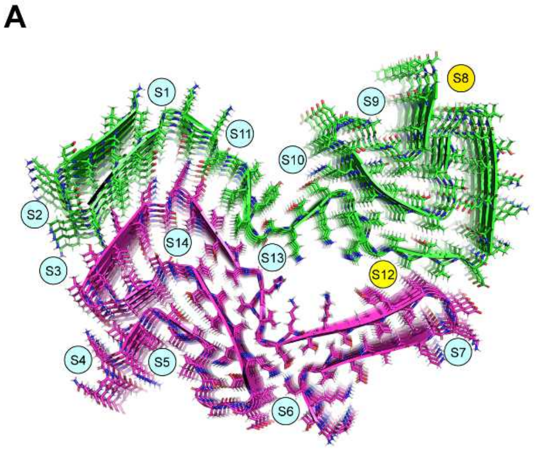


B

| | C05-05 | PM-PBB3 | PBB3 |
|---|---|---|---|
| S1 | -5.44 | -5.43 | -5.46 |
| S2 | -4.55 | -4.51 | -3.67 |
| S3 | -3.78 | -3.95 | -4.56 |
| S4 | -4.59 | -4.8 | -3.96 |
| S5 | -5.65 | -5.51 | -5.1 |
| S6 | -4.4 | -4.6 | -4.31 |
| S7 | -5.5 | -4.86 | -5.01 |
| S8 | -7.13 | -6.97 | -5.91 |
| S9 | -3.58 | -3.53 | -3.6 |
| S10-11 | -4.18 | -4.38 | -4.45 |
| S12 | -9.33 | -8.77 | -7.94 |
| S13 | -3.61 | -3.72 | -2.96 |
| S14 | -6.92 | -7.02 | -6.68 |

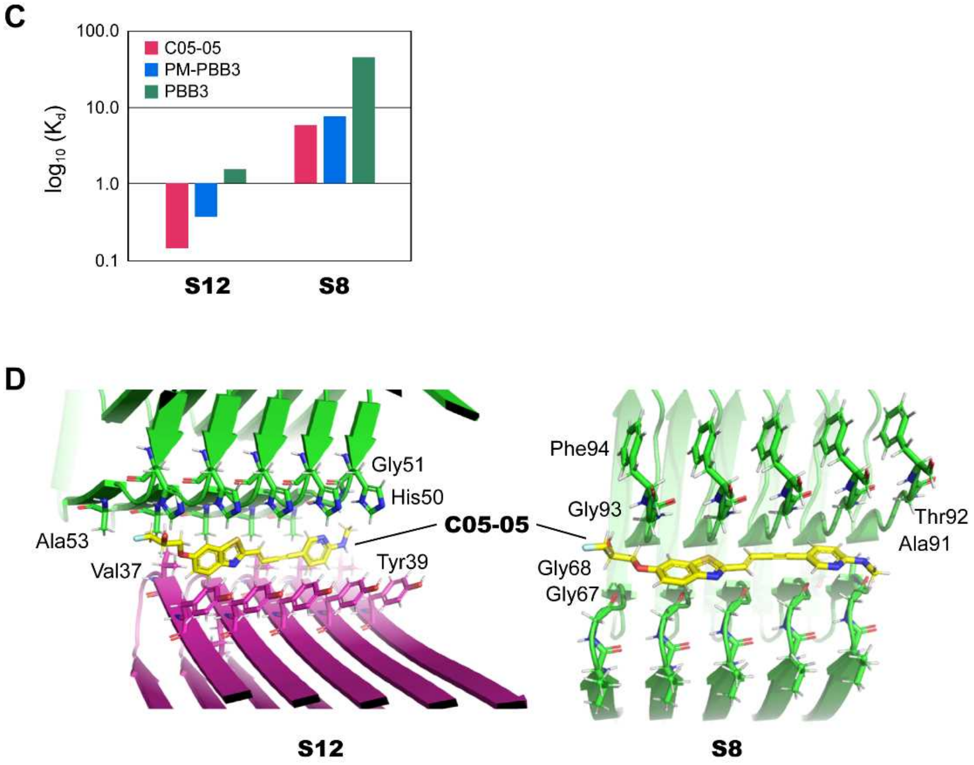


**Figure S5.** Binding mode of C05-05 in α-synuclein filaments predicted by molecular docking, Related to Figure 5.

(**A**) Docking sites of PET ligands to α-synuclein filaments explored by docking. PET ligands (C05-05, PM-PBB3, and PBB3) were docked to type I filaments of α-synuclein from MSA (PDB ID: 6XYO), and the binding affinity at each site (S1-S14) was calculated. Potential binding sites (S12 and S8) are shown in yellow, while the other sites with very weak compound binding are shown in sky blue. (**B**)

Binding scores (in kcal·mol$^{-1}$) of C05-05, PM-PBB3, and PBB3 in all the docking sites explored by docking. Docking site S12 shows the strongest binding, whereas S8 shows a moderate binding affinity for PET ligands. The PBB3 binds weakest, while C05-05 shows the strongest affinity. (**C**) Binding affinity (μM) of C05-05, PM-PBB3, and PBB3 for S8 and S12 calculated from docking energies. (**D**) Binding pose of C05-05 in S12 and S8. In S12, C05-05 is stacked between His50 and Tyr39 aromatic side chains mainly through hydrophobic interaction mediated by Val37, Ala53, and Tyr39. In S8, C05-05 forms hydrophobic interaction with Ala91 and polar interactions with Thr92. In both cases, the hydroxyl group on C05-05 forms a hydrogen bond with the backbone of Gly51 (S12) and Gly67-Gly68 (S8). Site S8 is identical in both type I and type II (PDB IDs: 6XYP and 6XYQ) filaments of α-synuclein from MSA, and therefore similar compound binding is seen. However, S12 is potentially absent in type II filaments as none of the ligands can dock into S12. The presence of Gly41 and Lys43 in the place of Val37 and Tyr39, respectively, changes the shape and electrostatic of S12.

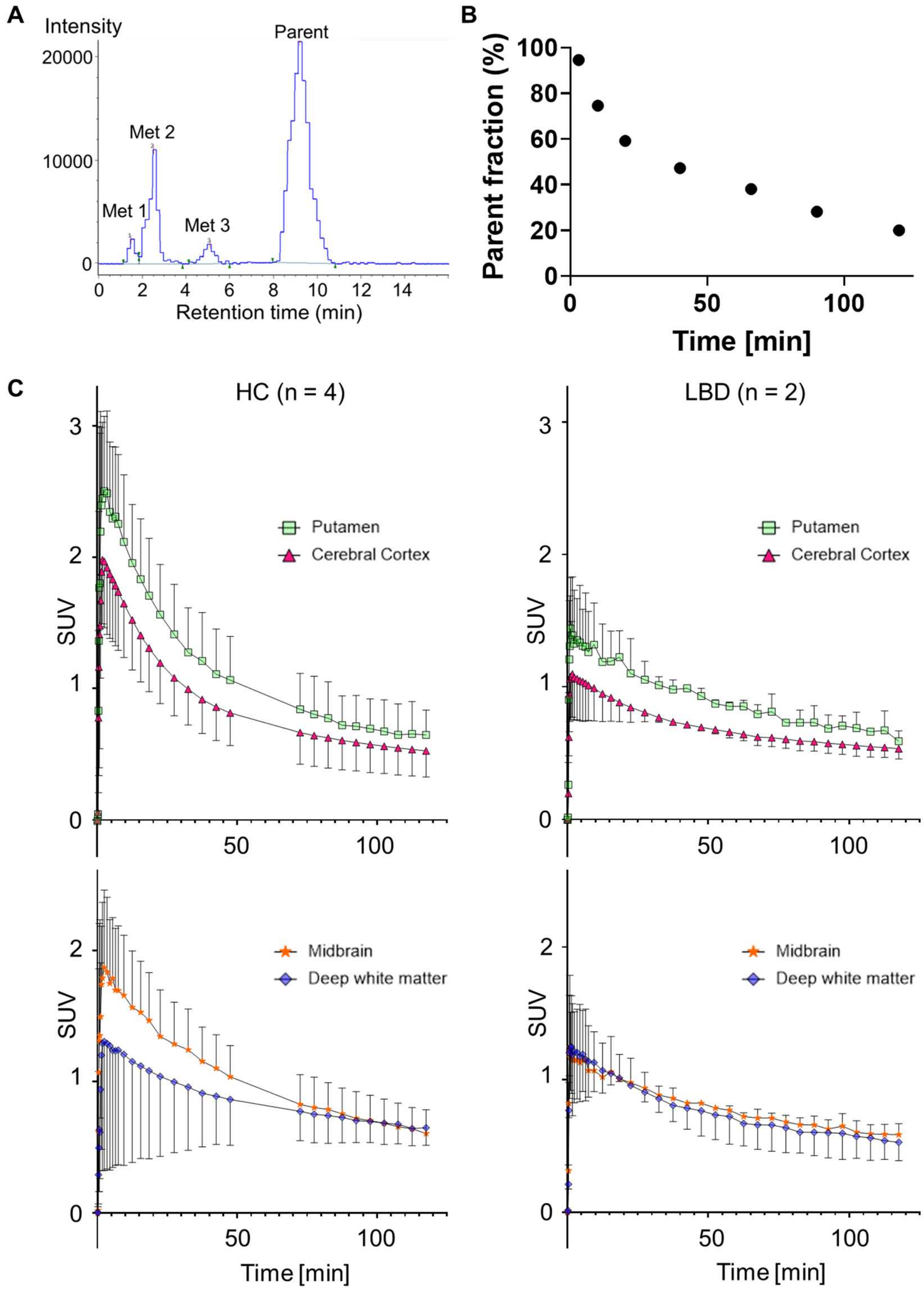


**Figure S6.** PET data in representative cases, Related to Figure 6.

(**A**) Radiochromatogram of plasma at 10 min after injection of $^{18}$F-C05-05. (**B**) Time course of parent fraction. (**C**) Time-radioactivity curve (TAC) for standardized uptake value (SUV) in the putamen and cerebral cortex (top row), and midbrain and deep white matter (bottom row) in healthy controls (HC) and patients with Lewy body disease (LDB).

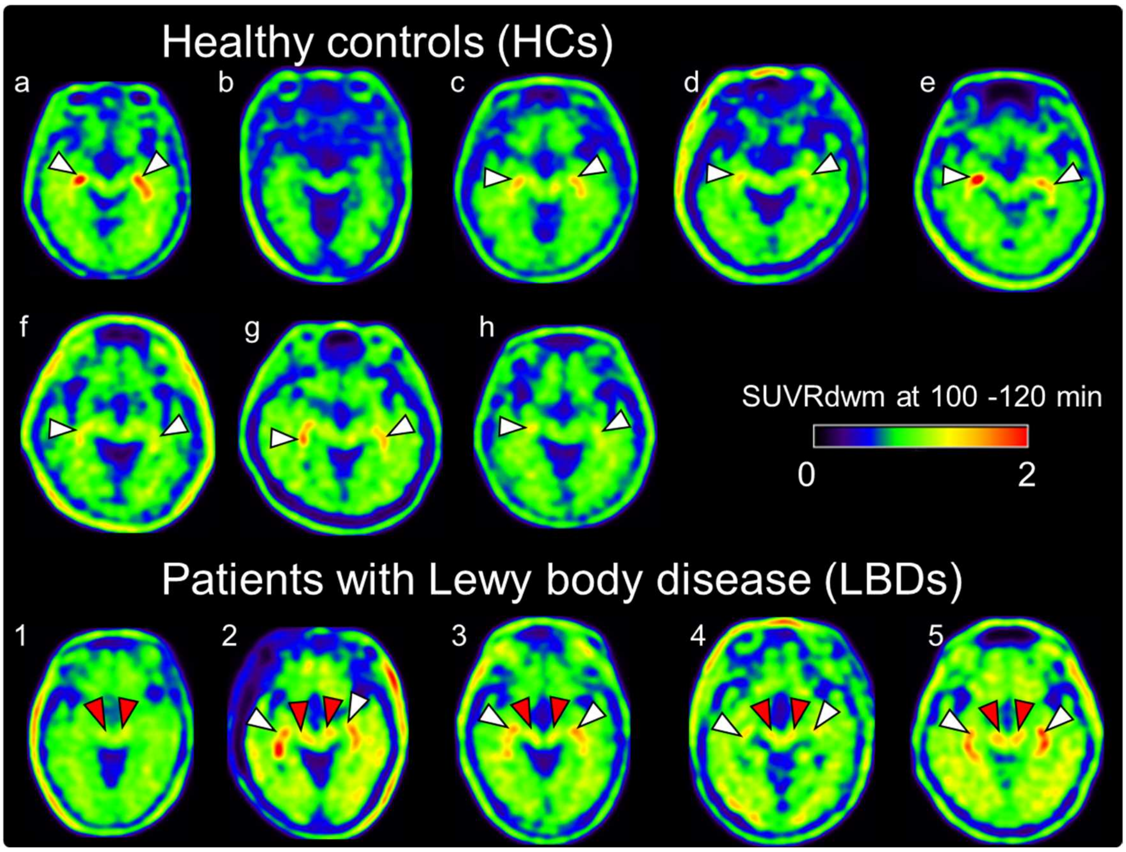


**Figure S7.** Parametric images showing SUVRs for $^{18}$F-C05-05 in a range from 0 to 2, Related to Figure 6.

PET images without superimposition on individual MRI data also show radioligand accumulations in the substantia nigra (red arrowheads). White arrowheads are considered nonspecific accumulation in the choroid plexus. The identity of the subjects are denoted by the same symbols (a – h and 1 – 5) as Figure 6.

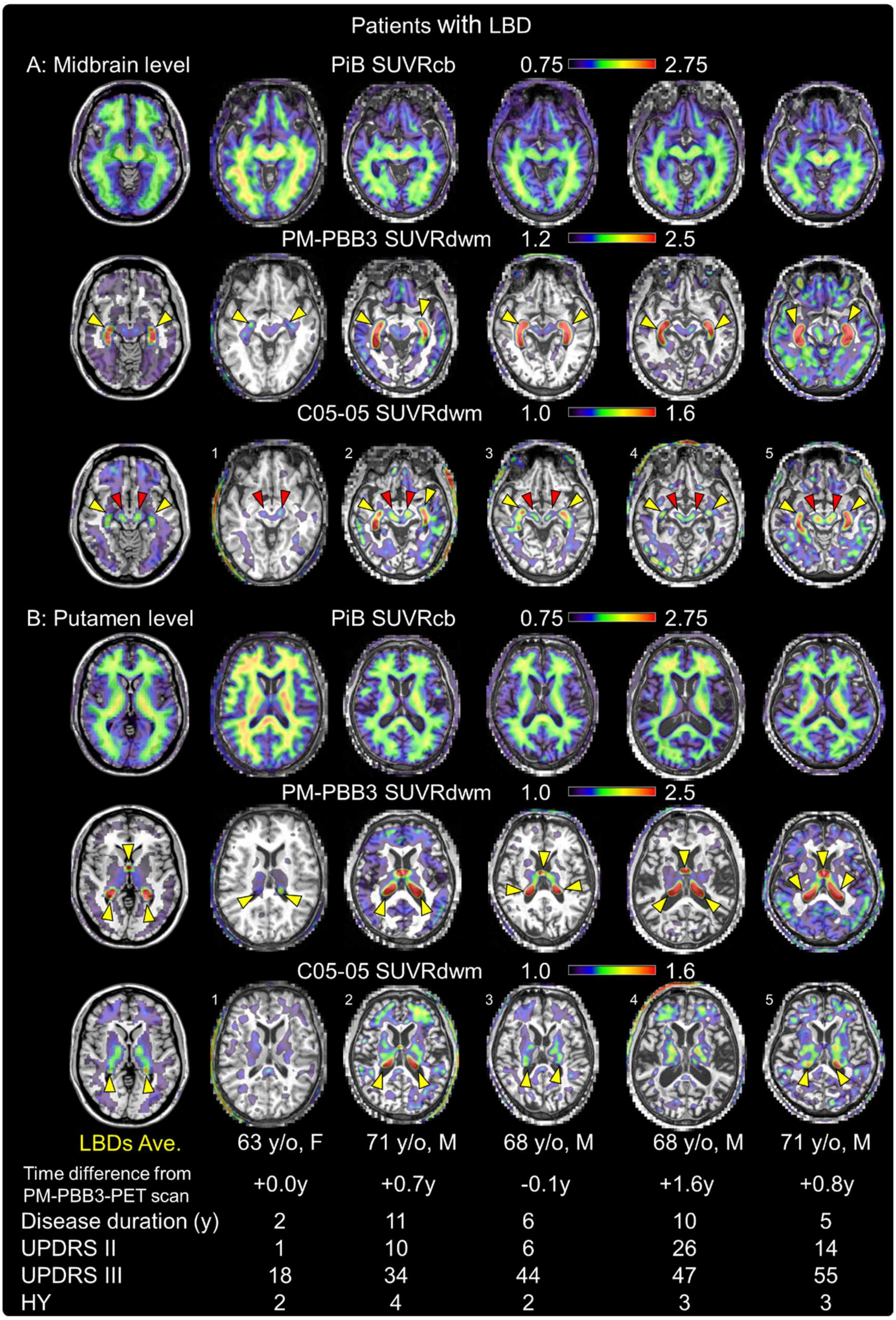


**Figure S8.** PET images with $^{11}$C-PiB, $^{18}$F-PM-PBB3, and $^{18}$F-C05-05 in the same individuals with Lewy body diseases (LBDs), Related to Figure 6.

Axial sections at the midbrain (**A**) and putamen (**B**) levels are displayed. The top, middle, and bottom rows in each panel show SUVR image for $^{11}$C-PiB, $^{18}$F-PM-PBB3, and $^{18}$F-C05-05, respectively, which were generated with the cerebellum gray matter (cb) or deep white matter (dwm) as reference regions. Age (years old; y/o) and sex (F, female; M, male) of the subjects are indicated, and images on the far left are averages of spatially normalized images derived from five LBD patients (LBDs Ave.) in the Montreal National Institute (MNI) space (midbrain level corresponding to Z = -11 and putamen level corresponding to Z = 1). In LBDs, there were accumulations of $^{18}$F-C05-05 radiosignals in the substantia nigra (red arrowheads) and nonspecific accumulations of $^{18}$F-PM-PBB3 and $^{18}$F-C05-05 radiosignals in the choroid plexus (yellow arrowheads). The identity of the subjects are denoted by the same symbols (1 – 5) as Figure 6. Abbreviations: HY (Hoehn and Yahr stage), UPDRS II (Unified Parkinson's Disease Rating Scale part II), UPDRS III (UPSRS part III), y (years)

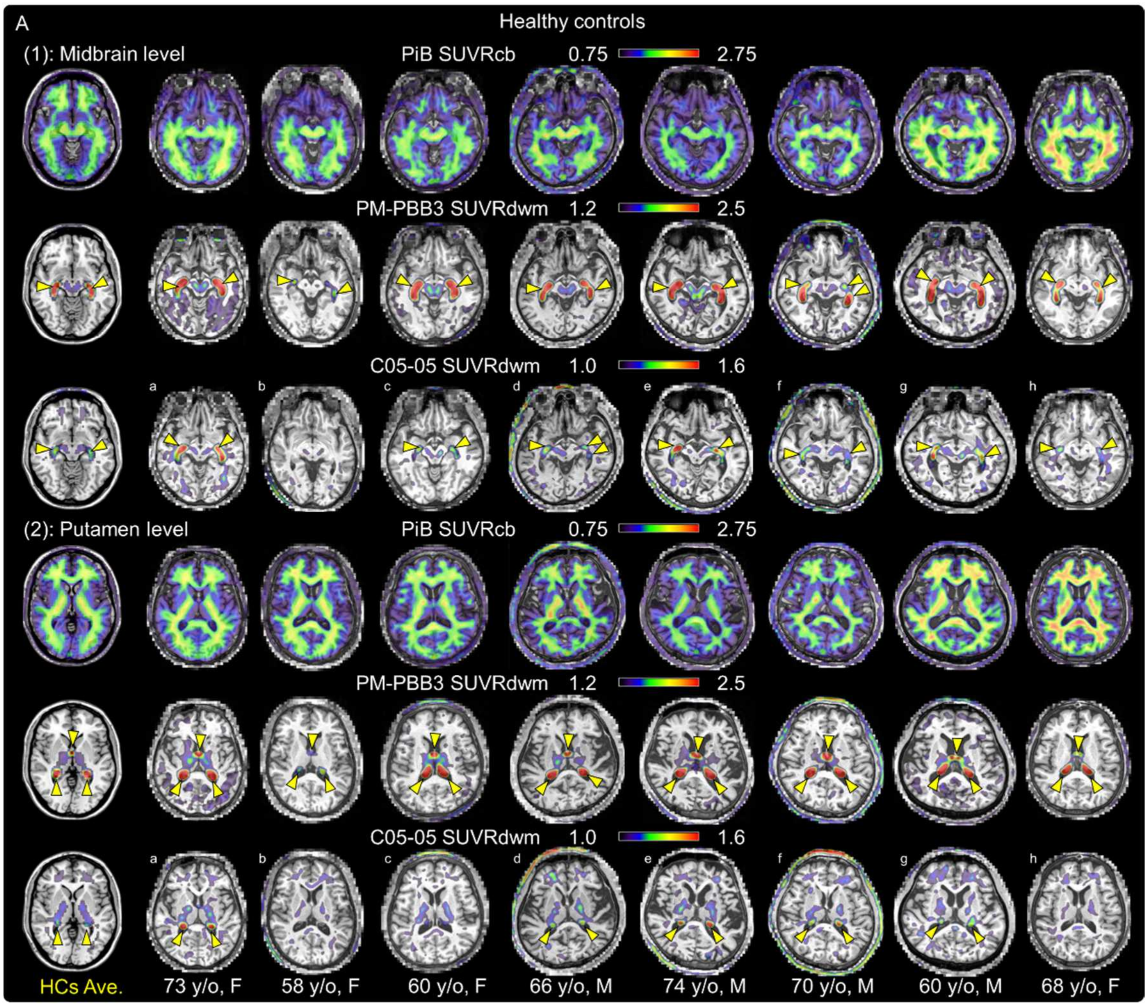
A
Healthy controls
(1): Midbrain level
PiB SUVRcb 0.75 2.75
PM-PBB3 SUVRdwm 1.2 2.5
C05-05 SUVRdwm 1.0 1.6
a
b
c
d
e
f
g
h
(2): Putamen level
PiB SUVRcb 0.75 2.75
PM-PBB3 SUVRdwm 1.2 2.5
C05-05 SUVRdwm 1.0 1.6
a
b
c
d
e
f
g
h
HCs Ave.
73 y/o, F
58 y/o, F
60 y/o, F
66 y/o, M
74 y/o, M
70 y/o, M
60 y/o, M
68 y/o, F

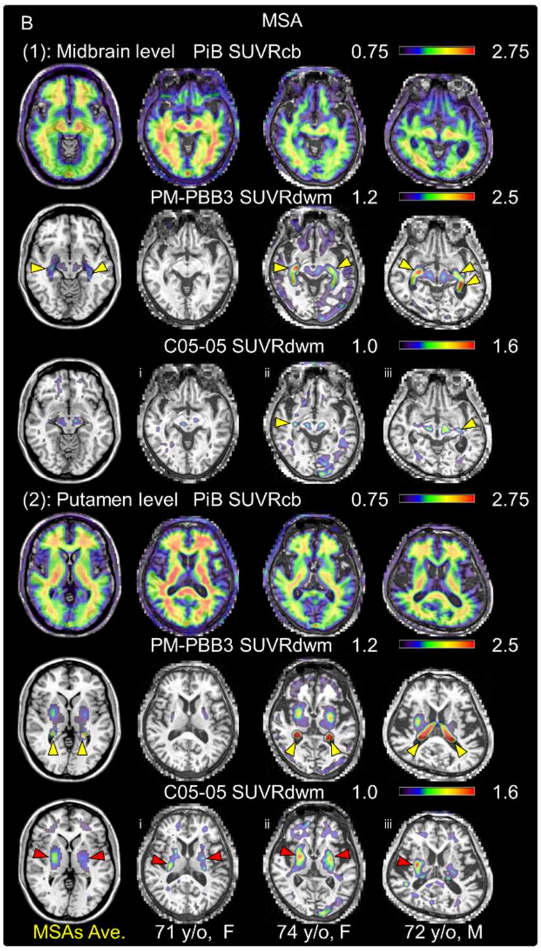


**Figure S9.** $^{11}$C-PiB and $^{18}$F-PM-PBB3 images of the same subject with $^{18}$F-C05-05 PET in healthy controls (HCs; **A**) and patients with multiple system atrophy (MSA; **B**), Related to Figure 6, 7.

Axial sections at the midbrain (**1**) and putamen (**2**) levels are displayed. The top, middle, and bottom rows in each panel show SUVR image for $^{11}$C-PiB, $^{18}$F-PM-PBB3, and $^{18}$F-C05-05, respectively, which were generated with the cerebellum gray matter (cb) or deep white matter (dwm) as reference regions. Age (years old; y/o) and sex (F, female; M, male) of the subjects are indicated, and images on the far left are averages of spatially normalized images derived from eight HCs (A) or three MSA patients (MSAs Ave.) in the Montreal National Institute (MNI) space (midbrain level corresponding to Z = -11 and putamen level corresponding to Z = 1). In MSA subjects, there were accumulations of $^{18}$F-C05-05 radiosignals in the lateral putamen (red arrowheads) and nonspecific accumulations of $^{18}$F-

PM-PBB3 and $^{18}$F-C05-05 radiosignals in the choroid plexus (yellow arrowheads). The identity of the HC (a – h) and MSA (i – iii) subjects are denoted by the same symbols as Figures 6 and 7, respectively.

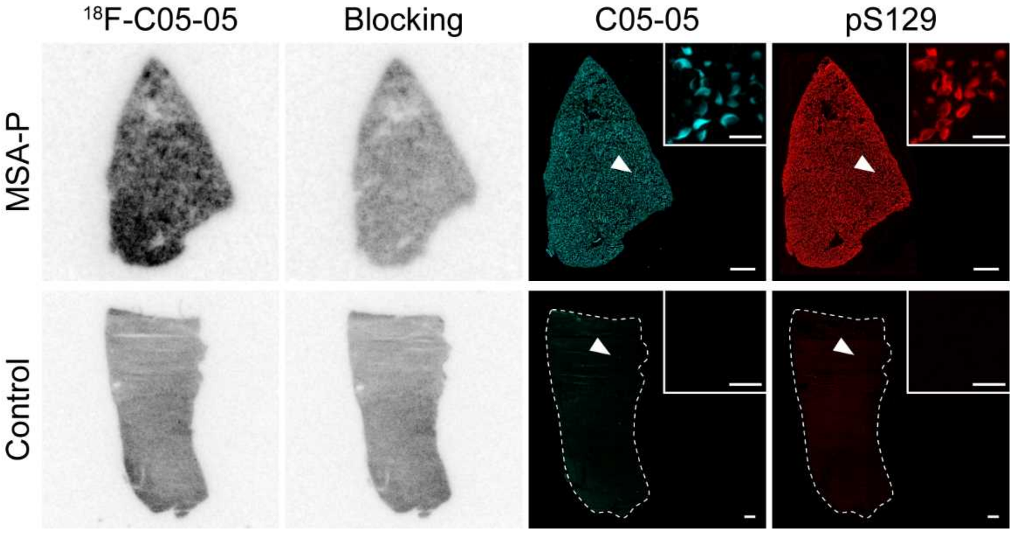


**Figure S10.** Autoradiographic labeling of putamen sections derived from an MSA-P with $^{18}$F-C05-05, Related to Figure 7.

Autoradiographic labeling of putamen sections derived from an MSA-P (top, also see Table S1) and a control (bottom, also see Table S1) cases with 5 nM of $^{18}$F-C05-05 in the absence (1st column from the left) and presence (2nd column) of 10 μM of non-radiolabeled C05-05. These sections were then used for fluorescence staining with non-radiolabeled C05-05 (30 μM) (3ed column) and pS129 (4th column), revealing abundant GCIs in the putamen of the MSA-P cases. Arrowheads indicate the location of the area magnified in the insets. Scale bars, 1 mm (low-power fields); 20 μm (insets).

**A**

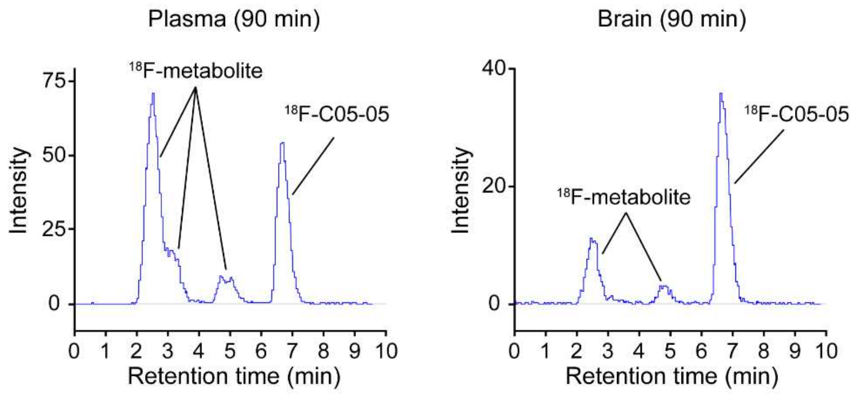


**B**

| | Plasma | | Brain | |
|---|---|---|---|---|
| Time (min) | Fraction (%) | | Fraction (%) | |
| | $^{18}$F-metabolite | $^{18}$F-C05-05 | $^{18}$F-metabolite | $^{18}$F-C05-05 |
| 5 | 7.6 | 92.4 | 0.9 | 99.1 |
| 30 | 44.4 | 55.6 | 11.8 | 88.2 |
| 60 | 57.1 | 42.9 | 19.8 | 80.2 |
| 90 | 65.5 | 34.5 | 28.2 | 71.8 |

**Figure S11.** Metabolite analysis of $^{18}$F-C05-05 in mice, Related to Figure 3.
(**A**) Radiochromatogram of plasma (left) and brain (right) at 90 min after intravenous injection of $^{18}$F-C05-05 in a representative case. (**B**) Unmetabolized $^{18}$F-C05-05 accounted for 92.4% and 99.1% of the total radioactivity in plasma and brain, respectively, at 5 min after intravenous injection, and 71.8% of the total radioactivity in the brain was derived from unmetabolized $^{18}$F-C05-05 even 90 min after intravenous injection.

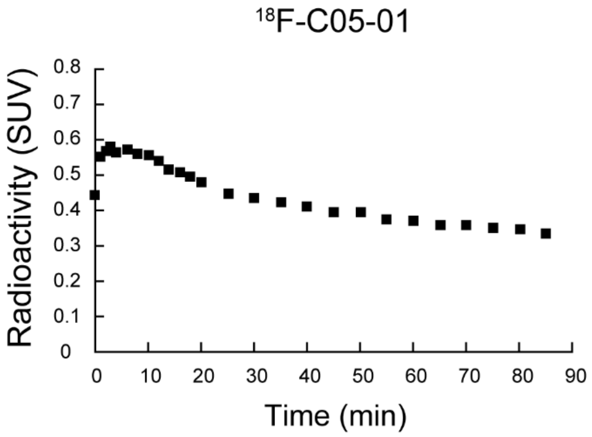


**Figure S12.** Time-radioactivity curve in the brain of wild-type mice in $^{18}$F-C05-01 PET, Related to Figure 3.

Time-radioactivity curves were generated in the frontal cortex of wild-type mice over 90 min after intravenous injection of $^{18}$F-C05-01. Data are presented as mean in two mice.

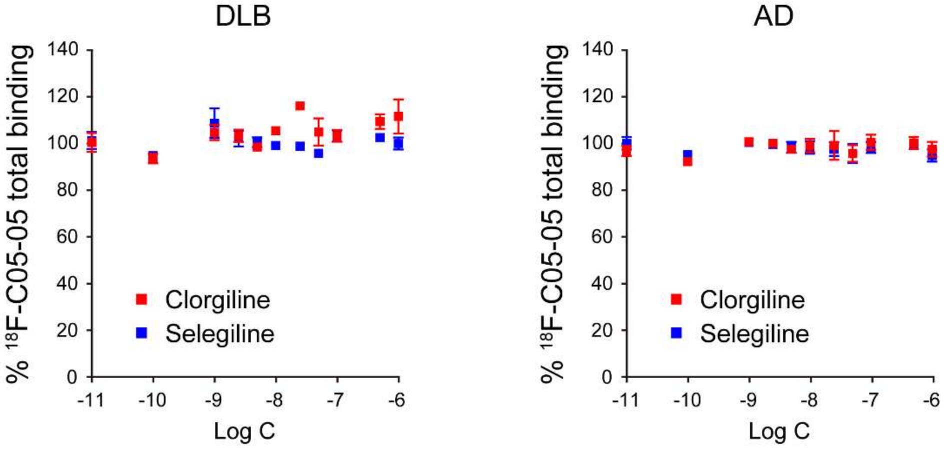


**Figure S13.** Inhibition of total binding of $^{18}$F-C05-05 by monoamine oxidase inhibitors in DLB and AD homogenates, Related to Figure 5.

Inhibition of total (specific + non-specific) binding of $^{18}$F-C05-05 by clorgiline (monoamine oxidase A inhibitor, red square) and selegiline (monoamine oxidase B inhibitor, blue square) in the DLB amygdala (left, also see Table S1) and AD frontal cortex (right, AD-2, also see Table S1) samples. Total binding of $^{18}$F-C05-05 was not inhibited by clorgiline and selegiline at varying concentrations, while Ki for clorgiline and selegiline was not determined due to failures of the model fitting. Data are mean ± SD in four samples and are expressed as % of the averaged total binding.

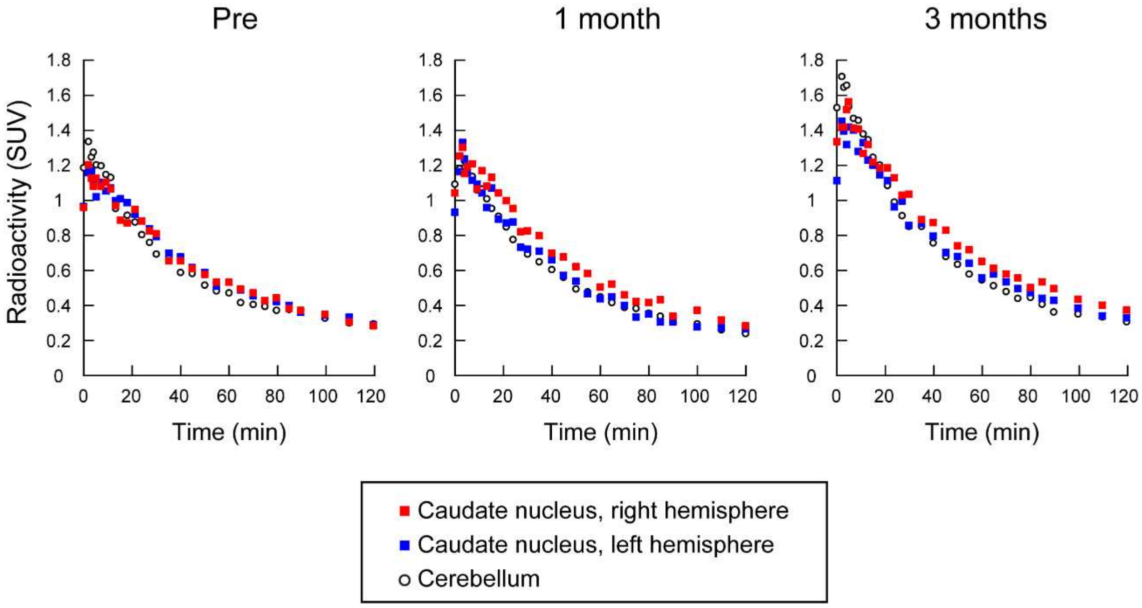


**Figure S14.** Time-radioactivity curves in the brain of an α-Syn marmoset in $^{18}$F-C05-05 PET, Related to Figure 4.

Time-radioactivity curves in the right and left caudate nuclei and cerebellum at 0 - 120 min after intravenous administration of $^{18}$F-C05-05 (89.6 ± 15.3 MBq) in an α-Syn marmoset before (Pre), and 1 and 3 months after inoculation.

**C05-03**

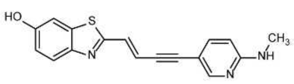

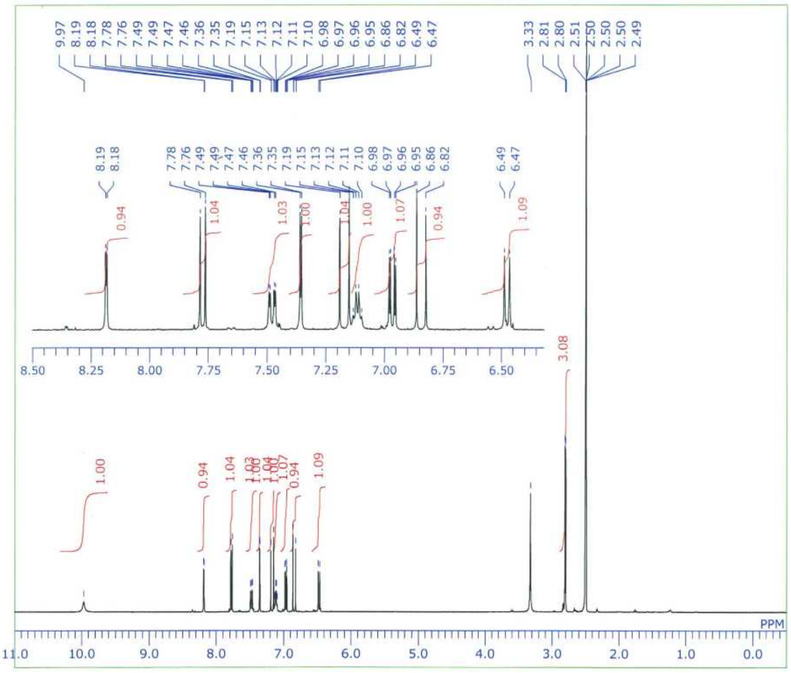

**C05-05**

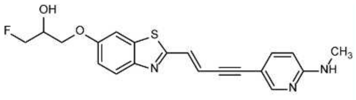

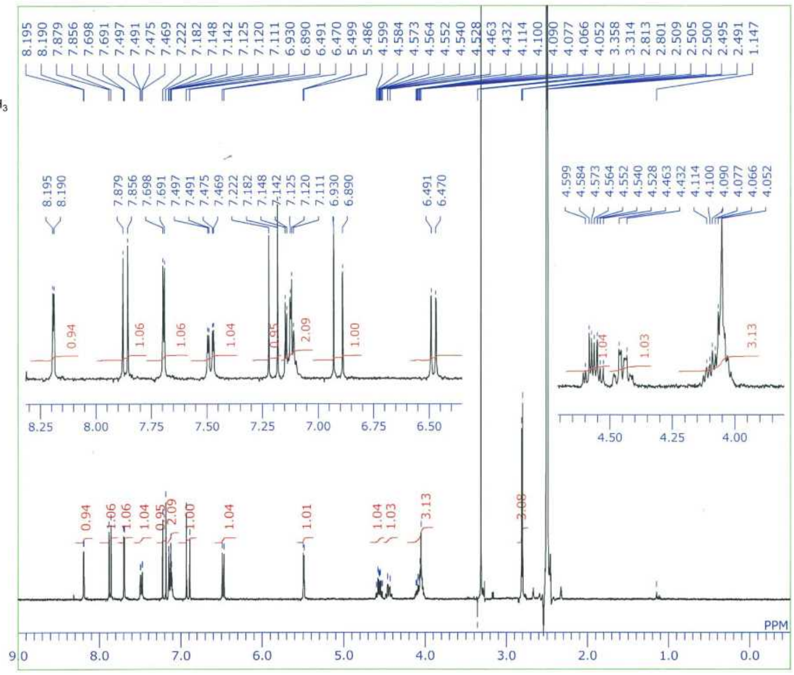

**Figure S15.** $^{1}$H NMR spectra for C05-03 and C05-05, Related to Figure 1.

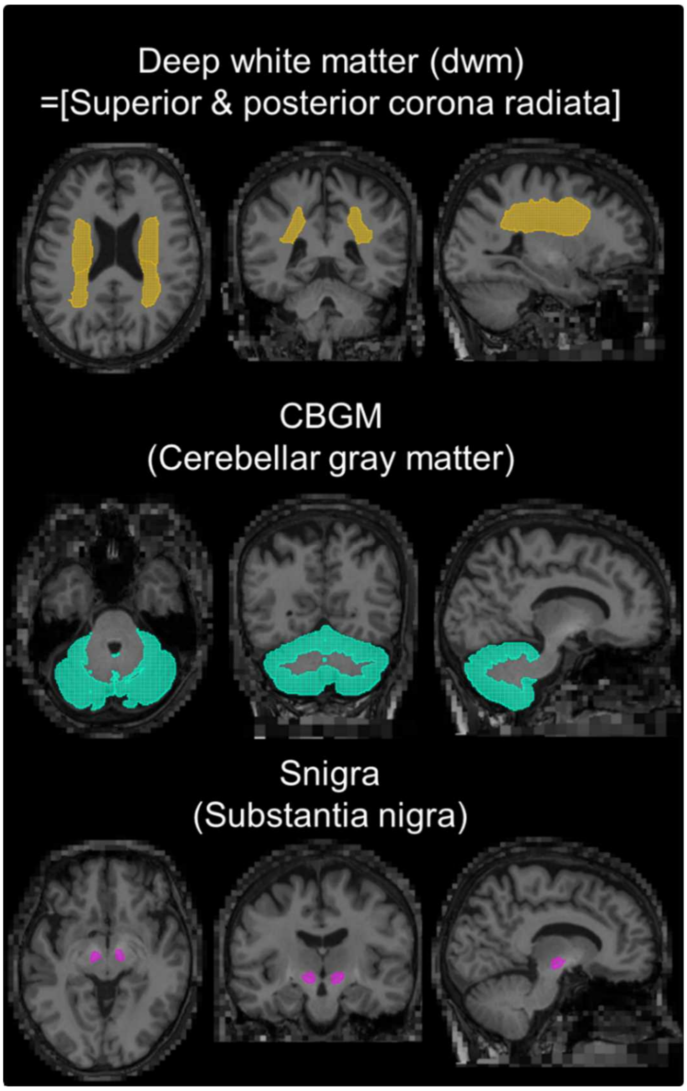


**Figure S16.** Representative reference and target region for PET SUVR, Related to Figure 6.

The SUVR image of C05-05 was obtained using dwm (yellow), while the SUVR images of PiB and PM-PBB3 were obtained using dwm and/or CBGM (light blue) as reference regions. Target regions were analyzed using black matter (pink). All regions of interest (ROIs) were obtained using M-Vision brain, an automatic brain segmentation software. The ROIs in the dwm were defined by combining superior corona radiata and posterior corona radiata from M-Vision brain. The volume (ccm) for all subjects was 0.54 ± 0.08 in the substantia nigra, 103.1 ± 13.1 in the CBGM, and 37.2 ± 4.5 in the dwm.

### SUPPLEMENTAL TABLE

| Case | Pathologic diagnosis | Clinical diagnosis | Age | Sex | Braak | CERAD score | Thal amyloid phase |
|---|---|---|---|---|---|---|---|
| DLB | TLBD | DLB or PSP | 57 | Male | I | | 0 |
| PDD | DDLB | PDD | 81 | Male | II | 0 | 1 |
| MSA-1 | MSA | Meniere disease | 82 | Male | II | | 1 |
| MSA-2 | MSA | MSA-P | 56 | Femal e | I | | 0 |
| MSA-3 | MSA | MSA-P | 59 | Male | I | | 1 |
| AD-1 | AD | FTLD-NOS | 69 | Male | III | 3 | |
| AD-2 | AD | AD probable | 58 | Male | III | 3 | |
| MSA-P | MSA | MSA-P | 62 | Femal e | I | 0 | 1 |
| Control | No significant alterations in the brain | POEMS syndrome | 49 | Femal e | I | 0 | 0 |

**Table S1.** Antemortem clinical diagnosis and neuropathologic description of postmortem human brain tissues, Related to Figure 1, 5, 6.
Braak, Braak neurofibrillary tangle stage; DLB, dementia with Lewy bodies; TLBD, Lewy body disease, transitional type; PSP, progressive supranuclear palsy; PDD, Parkinson’s disease with dementia; DDLB, Diffuse neocortical type of DLB; MSA, multiple system atrophy; MSA-P, MSA with predominant parkinsonism; AD, Alzheimer’s disease; FTLD-NOS, frontotemporal lobar degeneration not otherwise specified; POEMS syndrome, polyneuropathy, organomegaly, endocrinopathy, M-Protein, and skin changes syndrome.

| | HCs | LBDs | MSAs | *P* values |
|---|---|---|---|---|
| Number (race: all Asian) | 8 | 5 (PD = 4, DLB = 1) | 3 (MSA-P = 1, MSA-P with mild ataxia = 2) | |
| Age | 66.1 ± 6.2 | 67.8 ± 3.4 | 72.3 ± 1.5 | 0.09 |
| Female N [%] | 4 [50] | 1 [20] | 2 [67] | 0.38 |
| Amyloid-β positive | 0 | 0 | 0 | |
| Disease duration (year) | | 6.8 ± 3.7 | 2.3 ± 1.5 | 0.10 |
| UPDRS I | N.E. | 10.6 ± 6.3 | 10.0 ± 1.7 | 0.96 |
| UPDRS II | N.E. | 11.4 ± 9.5 | 23.3 ± 13.8 | 0.25 |
| UPDRS III | 1.6 ± 1.9 | 39.6 ± 14.2* | 62.0 ± 11.3** | <0.0001 |
| MMSE | 29.5 ± 0.8 | 29.6 ± 0.6 | 28.0 ± 1.7 | 0.15 |
| FAB | 16.9 ± 1.6 | 17.0 ± 1.2 | 14.3 ± 2.5 | 0.12 |
| GDS | 1.5 ± 1.5 | 5.2 ± 3.6 | 6.7 ± 3.5 | 0.04 |
| Apathy | 8.5 ± 5.7 | 14.8 ± 4.0 | 19.7 ± 12.5 | 0.15 |

*LBDs versus HCs post hoc Dunn’s test $p = 0.04$

**MSAs versus HCs post hoc Dunn's test $p = 0.004$

**Table S2.** Demographics of individuals with HCs, LBDs and MSAs included in the present study, Related to Table 1.

Abbreviations: FAB; Frontal Assessment Battery, GDS; Geriatric Depression Scale, HCs; Healthy Controls, LBDs; patients with Lewy body disease, MMSE; Mini-Mental State Examination, MSAs; patients with multiple system atrophy, MSA-P; MSA with predominant parkinsonism, N; number, N.E.; Not Examined, PD; Parkinson's Disease, DLB; Dementia with Lewy Body, UPDRS I, II, III; Unified Parkinson’s Disease Rating Scale part I, II and III.
